# Anxiety-Related Traits Are Associated with Subjective Biases but not Altered Threat–Safety Discrimination

**DOI:** 10.64898/2026.08.27.747474

**Authors:** Mana R. Ehlers, Hannah Stiffel, Alexandros Kastrinogiannis, Alina Koppold, Tina B. Lonsdorf

## Abstract

Anxiety-related traits (ARTs) have been linked to altered fear learning, but previous studies have typically examined different experimental phases and response systems, limiting the comparability of findings and the accumulation of consistent evidence. Here, we comprehensively examined associations between ARTs and fear conditioning across acquisition, extinction and renewal and across subjective, physiological and neural response systems in a well-powered sample (N = 267) using a two-day differential conditioning paradigm. ARTs were operationalized as a composite of trait anxiety, neuroticism, and intolerance of uncertainty and conditioned responding was assessed using skin conductance responses, fear-potentiated startle, US expectancy ratings, fear ratings, and functional magnetic resonance imaging. Higher ARTs were consistently associated with elevated subjective fear and US expectancy to both threat and safety cues during extinction and renewal, without corresponding elevations in physiological responding. At the same time, ARTs were not associated with threat–safety discrimination in subjective or physiological measures across phases, while neural associations were limited to reduced dorsal anterior cingulate cortex discrimination during early renewal. These findings suggest that ARTs are characterized by a CS unspecific cognitive bias toward heightened threat expectancy and evaluation rather than altered associative fear learning, highlighting the importance of distinguishing conditioned discrimination from general levels of responding across response systems.

## Introduction

Anxiety-and stress-related disorders affect approximately 28.8 % of the population (Kessler et al., 2005) and come with high individual and societal costs (Andlin-Sobocki et al., 2005; Konnopka & König, 2020). Alterations in fear learning and inhibition play a central role in the development and maintenance of such conditions (Duits et al., 2015; Kausche et al., 2025; Lissek et al., 2005). Fear learning processes show pronounced interindividual variability (Lonsdorf & Merz, 2017), which has been linked to anxiety-related traits (Eysenck, 1979; Haaker et al., 2015; Indovina et al., 2011; Lonsdorf & Merz, 2017; Morriss et al., 2015; Nelson & Shankman, 2011; Sjouwerman et al., 2020). Characterizing how such traits shape fear acquisition and extinction may help clarify mechanisms of vulnerability to anxiety-and stress-related disorders.

Fear learning can be investigated using differential fear conditioning, in which one initially neutral stimulus (CS+) is paired with an aversive unconditioned stimulus (US), whereas another stimulus (CS-) is not (Lonsdorf et al., 2017). CS discrimination, the response difference between the CS+ and CS-, is typically used as an indicator of conditioned threat responding. It can be assessed using subjective, physiological, and neurofunctional measures. During subsequent extinction training, the CS+ is presented without the US, resulting in a gradual reduction in conditioned responding through the formation of a new inhibitory memory (Myers & Davis, 2007). Conditioned responding may subsequently return when the CSs are presented in the acquisition context, a phenomenon known as renewal (Vervliet et al., 2013).

Anxiety-related traits – sometimes referred to as negative emotionality, dispositional negativity, or trait negative affect – reflect the tendency to experience negative affect more frequently, persistently, and intensely (Barlow et al., 2014; Chambers et al., 2004; Shackman et al., 2016). As established risk factors for anxiety and affective disorders, they are therefore prime candidates for investigating individual differences in fear learning (Barlow et al., 2014; Carleton et al., 2012; Chambers et al., 2004; Goldstein et al., 2018; McEvoy et al., 2019; McEvoy & Mahoney, 2012; Shackman et al., 2016).

In this study, ‘anxiety-related traits’ (ARTs) serves as an umbrella term encompassing three constructs selected in line with previous work (Sjouwerman et al., 2020): Trait anxiety, as assessed by the STAI-T (Version X; Spielberger et al., 1970), neuroticism, as assessed by the NEO-FFI-N (McCrae & Costa, 2004), and intolerance of uncertainty, as assessed by the IU-27 Scale (IUS; (Buhr & Dugas, 2002). All three constructs have been frequently examined in fear conditioning research (Lonsdorf & Merz, 2017; Sep et al., 2019). The instruments employed here represent well-established and widely used operationalizations of these constructs. In preparation for a meta-analysis, we identified the STAI-T and IUS as the most frequently investigated anxiety-related measures in the field (Bruntsch et al., 2024). Neuroticism, in turn, has often been used synonymously with the umbrella terms negative emotionality and dispositional negativity (Shackman et al., 2016). Their conceptual overlap is further reflected empirically, with particularly strong associations between the STAI-T and questionnaires assessing neuroticism (Klingelhöfer-Jens et al., 2025).

A growing number of studies have investigated associations between ARTs and conditioned fear responses across phases and outcome measures, revealing a rather heterogeneous body of evidence. Reported associations vary in direction, and null results are frequently reported (Bruntsch et al., 2024; Lonsdorf & Merz, 2017; see Supplementary Table 1 for details and references).

One potential source of this heterogeneity is variation in the outcome measures used to assess conditioned responding. Conditioned responding is assessed using multiple outcome measures, including physiological and behavioral measures (e.g., skin conductance response (SCR) and fear-potentiated startle (FPS)) and subjective ratings (e.g., US expectancy or fear ratings) each capturing partially distinct components of conditioned threat responding (Hamm & Vaitl, 1996; Lonsdorf et al., 2017; Ojala & Bach, 2020). Consequently, associations with ARTs may differ across response systems.

Among studies reporting trait-dependent differences, many support the notion of altered associative fear learning. During acquisition training, higher ARTs have been associated with reduced CS discrimination in SCR, FPS and US expectancy and fear ratings (Gazendam et al., 2013; Haddad et al., 2012; Morriss et al., 2019; Morriss, Macdonald, et al., 2016; Sjouwerman et al., 2020) and/or heightened CS-responses in FPS and fear ratings (Gazendam et al., 2013; Haaker et al., 2015; Sjouwerman et al., 2020). These patterns have been interpreted as indicating greater generalization across threat and safety cues and/or deficient safety learning. In line with this interpretation, during extinction, higher ARTs have been associated with heightened CS discrimination in SCR, FPS and US expectancy ratings (e.g., Morriss, 2019; Morriss, Christakou, et al., 2016; Morriss & van Reekum, 2019; Wake et al., 2020; Wroblewski et al., 2022) and/or heightened CS+ responses in SCR, US expectancy and fear ratings (Haaker et al., 2015; Morriss & van Reekum, 2019), indicating more persistent differential responding to threat and safety cues. These patterns, in turn, have frequently been interpreted as indicative of impaired extinction learning in individuals with higher ARTs.

Conversely, other studies have reported associations in the opposite direction. During acquisition training, higher ARTs have also been linked to greater CS discrimination in FPS and SCR (Chin et al., 2016; Indovina et al., 2011; Morriss et al., 2019), although responses to individual CS types were not reported, limiting interpretation of the underlying response pattern. Positive associations reported between ARTs and CS+ responses in SCR and fear ratings (Mertens et al., 2022; Wroblewski et al., 2022), may similarly reflect enhanced conditionability, as proposed previously in clinical populations (Lissek et al., 2005; Orr et al., 2000).

Several studies instead report elevated responding to both the CS+ and CS-within the same phase and outcome measure (i.e., fear ratings; Gazendam et al., 2013; Haaker et al., 2015; Haddad et al., 2012). Unlike changes in CS discrimination, such a general elevation cannot readily be attributed to altered associative learning and may instead reflect broader, non-associative differences. However, because these studies also reported associations consistent with altered CS discrimination or other associative effects (see above), their broader conclusions have predominantly focused on aberrant associative fear learning. Moreover, a large body of literature reports no associations between ARTs and conditioned responding across phases and outcome measures (e.g., Fredrikson & Georgiades, 1992; Morriss et al., 2020; Otto et al., 2007; Wendt & Morriss, 2022; see Supplementary Table 1).

Neuroimaging studies suggest that ARTs – despite a heterogeneous body of evidence and null findings (Wendt & Morriss, 2022; Wroblewski et al., 2022) – may be associated with altered neural responses during fear learning. During acquisition training, higher ART levels have been linked to stronger engagement of brain regions known to be involved in threat and safety processing and associative learning. For example, they have been related to greater CS discrimination in the amygdala, putamen, and thalamus (Sjouwerman et al., 2020), and to stronger amygdala responses to predictive versus safe cues (Indovina et al., 2011). Higher levels have also been linked to greater engagement of prefrontal regions such as increased activity in medial and dorsomedial rostral prefrontal cortex in a threat of shock task (Morriss et al., 2022). In addition, they have been linked to stronger functional connectivity between the amygdala and hippocampus as well as between the amygdala and prefrontal regions during acquisition training (Tzschoppe et al., 2014). During extinction, evidence similarly remains mixed with null results (Wendt & Morriss, 2022), and studies suggesting that higher ART levels are associated with greater amygdala and vmPFC activity in response to threat relative to safety cues during late extinction (Morriss et al., 2015), potentially reflecting persistent threat appraisal alongside increased recruitment of prefrontal processes involved in safety signaling and updating threat value. In contrast, during early extinction training individuals with higher ART levels show enhanced CS discrimination in the vlPFC, driven by more pronounced deactivation to the CS+ (Wroblewski et al., 2022). Beyond acquisition and extinction training, neuroimaging evidence is scarce, particularly with regard to renewal. One study reported that higher ART levels were associated with greater neural responses to safety relative to threat cues during reinstatement test and re-extinction phase in regions of the fear network, including the thalamus and anterior insula (Wroblewski et al., 2022). Together with persistent physiological CS discrimination observed during extinction, these neural response patterns were interpreted as reflecting reduced adaptation to changing threat contingencies and continued responding based on previously acquired threat associations in individuals with higher ART levels.

A central challenge for cumulative knowledge building is a substantial fragmentation of the literature, which hampers the integration of findings across studies. Individual studies typically focus on different and often narrow combinations of outcome measures, ART constructs and experimental phases, making it difficult to determine whether divergent findings reflect genuine differences across response systems, traits, stimuli and phases or methodological differences. Results from preceding experimental phases are not always reported, further limiting interpretability. Furthermore, studies often rely on small samples (see Lonsdorf et al. (2017) for a review) limiting statistical power to reliably detect individual differences (Mar et al., 2013; Schönbrodt & Perugini, 2013). For an overview of previous studies, see Supplementary Table 1 and 2.

Against this background, the present study investigated associations between ARTs and fear conditioning across acquisition, extinction, and renewal, using multimodal outcome measures (i.e., SCR, FPS, US expectancy and fear ratings, and fMRI) in a single well-powered sample. By providing the most comprehensive investigation to date, the present study seeks to contribute to the existing literature by allowing for a more fine-grained and comprehensive understanding beyond the current evidence base.

Based on previous reports of ART-related differences in associative fear learning, we expected ARTs, operationalized as a composite score of trait anxiety, neuroticism, and intolerance of uncertainty, to be associated with CS discrimination across acquisition, extinction, and renewal. Given the heterogeneity of previous findings, particularly in the direction of associations, no directional hypotheses were specified. Responses to the CS+ and CS-were additionally examined separately to characterize the response patterns underlying potential differences in CS discrimination.

## Materials and Methods

### Participants

The current study was part of a larger project where participants completed in addition to a differential two-day fear conditioning paradigm, an approach-avoidance task in virtual reality and a freezing related task on two consecutive days (Kastrinogiannis et al., 2025; Koppold et al., 2026).

A total of N = 277 participants were recruited via online advertisements and bulletin boards at the University Medical Center Hamburg-Eppendorf (UKE). 16 participants completed the task in the behavioral lab, the remaining participants performed the study in the scanner. In addition to a phone screening, all participants who completed the task in the MR scanner underwent an online or in person screening with a medical doctor to ensure inclusion criteria for participation in the current study in general and in MR studies in particular were met. Participants were required to be between 18 and 50 years old and right-handed. Exclusion criteria included participation in medication studies, regular medication intake (with the exception of oral contraceptives, thyroid or allergy medications, or occasional use of over-the-counter drugs), pregnancy or breastfeeding, as well as MRI-specific contraindications (e.g., claustrophobia or non-MRI-compatible metal implants or objects such as copper intrauterine devices).

Of the 277 recruited participants, eight were excluded because they completed an earlier version of the paradigm (*n* = 4), discontinued the study (*n* = 3), or were not assigned a participant ID (*n* = 1). Two additional participants were excluded because, owing to a technical error, they did not receive any US presentations during acquisition, resulting in a final sample of *N* = 267 participants (mean age = 26.6 years, *SD* = 5.3, range = 18 - 50 years; 170 female, 96 male, and 1 diverse). Sample sizes varied across outcome measures because of procedural or technical problems and insufficient data quality, particularly for SCR and FPS recordings obtained in the MR scanner. Sample sizes varied across outcome measures because of missing recordings and measure-specific data-quality exclusions. Participants were retained for all analyses for which valid data were available. The exclusion criteria and final sample size for each outcome measure are reported in the Supplementary Material.

All participants provided written informed consent in accordance with the Declaration of Helsinki and received financial compensation of 80 euros. The study was approved by the Ethical Review Board of the General Medical Council Hamburg (PV5808).

### Questionnaires

All participants filled in a set of questionnaires, including the State-Trait Anxiety Inventory (STAI-T; Grimm, 2009; Spielberger et al., 1970), the neuroticism subscale of the NEO-Five-Factor Inventory (NEO-FFI-N; Gerhard, 1999; McCrae & Costa, 2004) and the Intolerance of Uncertainty Scale (IUS; Buhr & Dugas, 2002; Gerlach et al., 2008). All scores were z-transformed. For each participant, z-scores were then combined to determine a global composite score of anxiety-related traits (ARTs).

#### Trait anxiety

The trait scale of the State-Trait Anxiety Inventory (STAI-T; Grimm, 2009; Spielberger et al., 1970) comprises 20 self-reported items (e.g. “I feel that difficulties are piling up so that I cannot overcome them“), answered on a 4-point Likert scale ranging from 1 (almost never) to 4 (almost always), resulting in sum scores between 20 and 80. High scores indicate a predisposition to perceive different situations as threatening or stressful, and to react anxiously. However, it should be noted that the STAI-T has been shown to not tap specifically into anxiety but also other dimensions of negative emotionality – most notably depressive symptoms (Bados et al., 2010; Balsamo et al., 2013; Bieling et al., 1998; Klingelhöfer-Jens et al., 2025; Knowles & Olatunji, 2020).

#### Neuroticism

Neuroticism as a trait disposition was assessed by using the neuroticism subscale of the NEO-Five-Factor Inventory (NEO-FFI-N; Gerhard, 1999; McCrae & Costa, 2004), which consists of 12 self-reported items (e.g. “I often feel tense and jittery“). Items are rated on a 5-point Likert scale ranging from 0 (strongly disagree) to 4 (strongly agree) and allowing participants to score between 0 and 48 points. High scores indicate a tendency to experience and express negative emotions (Ormel et al., 2013).

#### Intolerance of uncertainty

The Intolerance of Uncertainty Scale (IUS; Buhr & Dugas, 2002; Gerlach et al., 2008) was applied. The scale includes 27 self-reported items (e.g. “I can’t stand being undecided about my future”), which are evaluated on a 5-point Likert scale ranging from 1 (not at all characteristic of me) to 5 (entirely characteristic of me) and allowing sum scores between 27 and 135. Participants with high scores tend to experience uncertain future events as stressful and threatening (Freeston et al., 1994).

### Stimuli

#### Electrotactile stimulus

The US, an electrotactile stimulus, was administered to the back of the participant’s right hand between the metacarpal bones of the index and middle finger through a 1 cm diameter platinum pin surface electrode (Specialty Developments, Bexley, UK). The stimulation consisted of three 2ms rectangular pulses with an interpulse interval of 50 ms. The pulse was delivered via a Digitimer DS7A constant current stimulator (Welwyn Garden City, Hertfordshire, UK). US intensity was calibrated individually to be clearly unpleasant but not painful (aiming for an unpleasantness rating of 7-8 out of 10).

#### Visual stimuli

Two black snowflakes, presented on yellow or blue background (context A or context B, respectively), served as CS+ and CS-. The allocation of stimuli, stimulus order and background colors were counterbalanced across participants.

#### Startle probe

A burst of 95 dB white noise was used to elicit the startle eyeblink reflex. The sound was presented via MR-compatible earphones (Sensimetrics Corporation, Gloucester, MA, USA) in the scanner or via headphones (Sennheiser, Wedemark, Germany) in the behavioral lab.

### Experimental Design

Laboratory assessments were conducted on two consecutive days, with habituation and fear acquisition training on day one and extinction training followed by a ROF test (i.e., renewal) on day two (24 h later). All experimental phases were performed in the MR scanner for the majority of participants (see above). All participants started with an individual shock calibration. This was followed by a habituation phase. The instructed fear acquisition training began after participants were informed of the CS+/US contingency, but not of the reinforcement rate. On day two, delayed extinction training took place. Afterwards, a return of fear test (i.e., renewal) was conducted following an ABA paradigm. Renewal was completed in context A, the same context as in the acquisition, while extinction was performed in context B. For additional procedural details see Figure 1.

**Figure 1.**
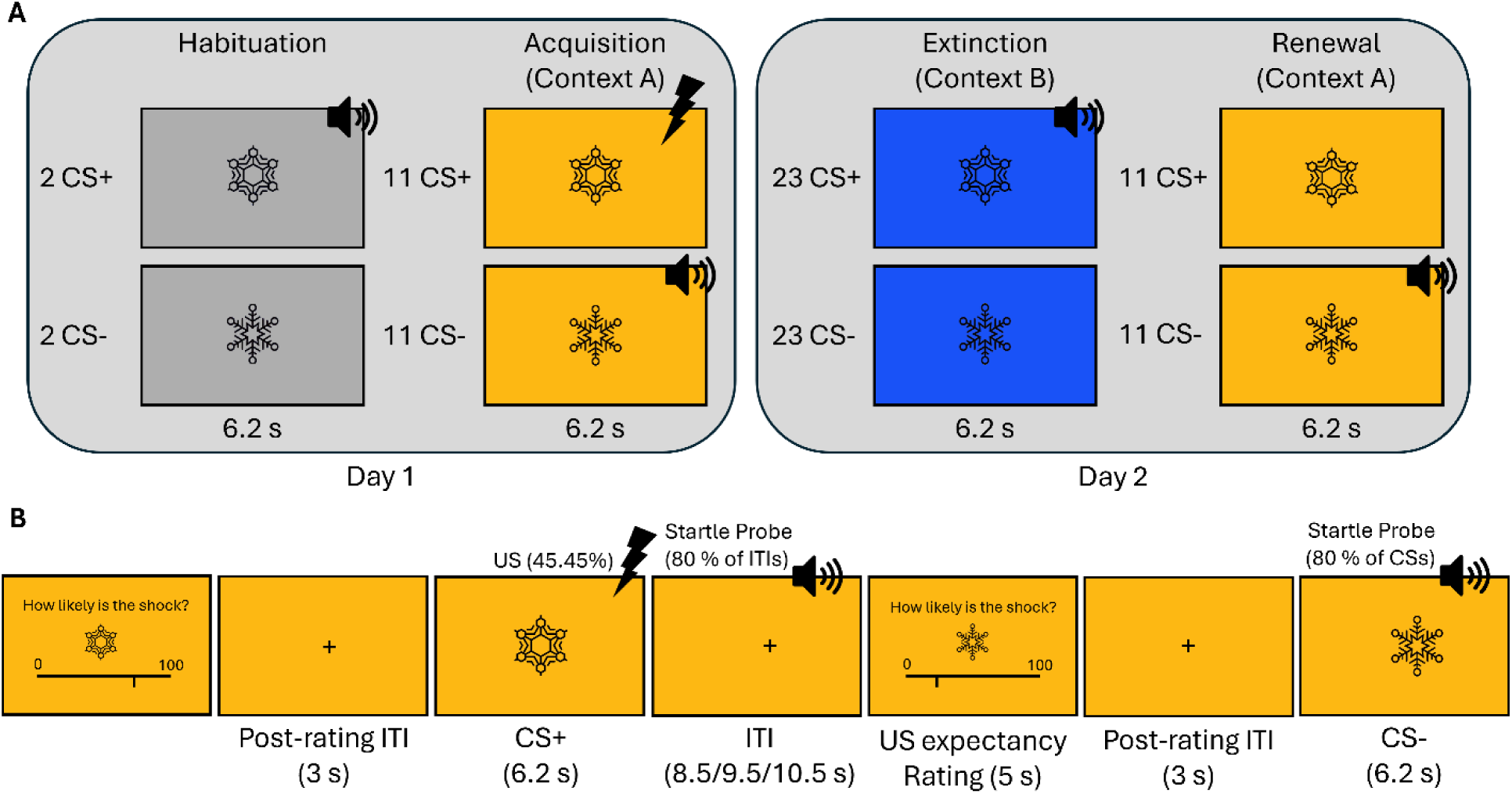
Experimental design. (A) Overview of the experimental procedure across two days. On day 1, participants completed habituation followed by instructed fear acquisition training in context A. During habituation, CS+ and CS-were each presented twice. During acquisition training, CS+ and CS-were each presented 11 times; the CS+ was partially reinforced (5/11 trials, 45.45%) with the US delivered shortly (i.e., 160 ms) before stimulus offset, whereas the CS-was never reinforced. Acquisition training was fully instructed. On day 2 (∼24 hours later), participants underwent extinction training in a different context (context B), in which CS+ and CS-were each presented 23 times. This was immediately followed by a renewal test in the original acquisition context (context A), where CS+ and CS-were again each presented 11 times. The US was only presented during acquisition training. **(B) Example trial structure.** Each trial began with an US expectancy rating, followed by a fixation period (3 s) and CS presentation (6.2 s). On reinforced acquisition trials, the US occurred 100 ms before CS+ offset. Startle probes were delivered on 80% of CS presentations (jittered at 4.5–5 s after CS onset) and during inter-trial intervals (ITIs; 8.5– 10.5 s). Trial timing and probe onset were counterbalanced across participants.

### Fear and Expectancy Ratings

#### Fear ratings

Fear ratings were completed before and after acquisition, extinction and renewal. Participants had to indicate on a 100-point visual analog scale (VAS) how much “stress, fear and tension” they experience when they see the CS+ and CS-respectively. Seven seconds response time was allocated to answer on a range from zero (answer = none) to 100 (answer = a lot).

#### US expectancy ratings

US expectancy ratings were collected before every CS-block. Participants used a VAS scale ranging from 0 to 100 % to indicate how high they estimate the likelihood that the US will be delivered the next time the CS+ or CS-are presented. If no response was given and confirmed within the 5 second time window, it was counted as missing.

### Physiological Measurements

#### Skin conductance response

SCR data were collected with a Biopac MP100-amplifier system (BIOPAC Systems Inc, Goleta, California, USA) and Spike 2 software in the MR scanner (Cambridge Electronic Design, Cambridge, UK) and Acqknowledge 3.9.2 in the behavioral lab. Two self-adhesive Ag/AgCl electrodes were filled with isotonic electrode gel and placed on the distal and proximal hypothenar on the palmar side of the left hand. Data were recorded throughout the experiment with a sampling rate of 1000 Hz, a gain of 5 µΩ and was down-sampled to 10 Hz during preprocessing.

Data were scored semi-manually using the trough-to-peak (TTP) method implemented in the custom-made program EDA view (developed by Prof. Dr. Matthias Gamer, University of Würzburg). Responses were scored when the response onset occurred in a time window of 0.9 to 4.5 s after stimulus onset with a rise time of max. 5 s (Boucsein, 2012; Sjouwerman & Lonsdorf, 2019). Each response was checked visually and the algorithm-suggested response corrected if necessary. Trials with artifacts or excessive baseline activity were scored as missing values. Response amplitudes smaller than 0.01 µS were set to zero (Lonsdorf et al., 2019). In order to account for individual differences, raw SCR data were log transformed and range corrected by dividing each response by the maximum (to CS or US) for each participant and day. Participants who had more than 2/3 missing or zero response to the US were classified as “non-responders” (n = 54) and were excluded from the analysis (Lonsdorf et al., 2019).

#### Fear potentiated startle

In the MR scanner, startle-eyeblink response data recording through electromyography followed a previously established protocol (Kuhn et al., 2020) using BrainAmp ExG MR amplifier including a SyncBox device for synchronization of recorded data and MR gradient switching (Mandelkow et al., 2006) and BrainVision Recorder software (Brain Products GmbH, Gilching, Germany). For electrode placement, a FaceEMG Cap-MR (EasyCap GmbH, Herrsching, Germany) with five Ag/AgCl electrodes was used to ensure safe wire management and facilitate artifact reduction within the MR environment. The skin under the participant’s left eye for placement on the orbicularis oculi muscle was prepared with abrasive electrode gel. One additional electrode was placed on the participant’s forehead and two electrocardiogram electrodes at the back. The resistance threshold for all five electrodes was kept under 20 kOhm. Data were recorded with a sampling rate of 5000 Hz with a signal resolution of 0.1 μV within a frequency band of 0.016 and 250 Hz. Data preprocessing performed with BrainVision Analyzer software followed previous recommendations (Kuhn et al., 2020; Lindner et al., 2015) and included MR gradient correction by subtracting an artifact template from the raw EMG that was created from artifacts averaged over seven echo-planar-imaging (EPI) volumes. The data were down-sampled to 1000 Hz, eye movement related activity was reduced by applying a low pass filter with a cutoff of 60 Hz, with a time constant of 0.0027 and 48 db/oct and data were rectified before manual scoring.

In the behavioral lab, FPS data were collected with a Biopac MP100-amplifier system (BIOPAC Systems Inc, Goleta, California, USA) and Acqknowledge 3.9.2 software. The skin was prepared with abrasive electrode gel and three Ag/AgCl were attached on the participant’s orbicularis oculi muscle and on their forehead. Data were recorded with a sampling rate of 1000 Hz and a gain of 5000 and were band pass filtered online (28 – 500 Hz). The data were integrated (averaged over 20 samples) and rectified.

Data from both the MR environment and the behavioral lab were scored semi-manually as foot to peak (within a time-window 20 – 150 ms post startle probe onset) using EDA view and following published guidelines (Blumenthal et al., 2005). If a response was confounded by a blink in the period up to 50 ms before startle probe onset, the response was counted as missing. Each participant’s raw scores were t-transformed within each experimental phase.

### Statistical Analysis of Behavioral and Physiological Data

In order to assess the success of the experimental manipulation, paired *t*-tests were performed to compare CS+ and CS-responses during acquisition training, extinction training, and renewal, using responses averaged across all trials for SCR, FPS, and US expectancy ratings, and post-phase ratings for fear ratings. In addition, separate 2 × 2 ANOVAs with the factors stimulus (CS+ vs. CS-) and time (first vs. second half of the phase) were conducted to assess potential changes in CS discrimination across the first and second halves of extinction and renewal.

As the three questionnaires assess overlapping facets of broadly ‘anxiety-related traits’ and were therefore combined into a standardized composite index. For each scale, a participant’s sum score was z-transformed (relative to mean and standard deviation of the whole sample). For each participant, a combined score was derived by averaging across the z-scores from the three questionnaires (Abend et al., 2019, 2022). For the two physiological outcome measures and the ratings, CS discrimination indices were derived by subtracting CS-responses from CS+ responses. Correlation analyses were conducted separately for the outcome measures SCR, FPS, expectancy and fear ratings, using CS+, CS-, and CS discrimination values. To control for multiple comparisons, the Benjamini-Hochberg procedure was applied. For BOLD fMRI data, correlations between anxiety-related traits and the contrast CS+ > CS-were computed and assessed using small volume correction in a range of ROIs.

Statistical analyses and visualization were performed with R version 4.6.1 (2026-06-24) using the packages cowplot, dplyr, ggcorrplot, ggplot2, ggpmisc, ggpubr, gridExtra, patchwork, psych, purrr, rstatix, tidyr. All analysis code is available at https://github.com/manaehlers/ART-fear-conditioning.

### Functional magnetic resonance imaging (fMRI)

#### Data acquisition

Functional MRI data were collected on a 3 Tesla PRISMA whole-body scanner (Siemens Medical Solutions, Erlangen, Germany) equipped with a 64-channel head coil, using an echo planar imaging (EPI) sequence (repetition time = 1910 ms, echo time = 30 ms, 54 slices, slice thickness = 1.7 mm with 1 mm gap, field of view = 224 × 224 mm). High-resolution T1-weighted anatomical scans were obtained via a magnetization-prepared rapid gradient echo (MPRAGE) sequence (TR = 7.1 ms, TE = 2.98 ms, 240 slices, slice thickness = 1 mm, field of view = 192 × 256 mm).

#### Regions of interest

Based on prior research on neural markers of fear conditioning (Bechara et al., 1995) and meta-analytic findings (Fullana et al., 2016, 2018), we defined 11 regions of interest (ROIs): amygdala, hippocampus, caudate nucleus, putamen, pallidum, ventral striatum, thalamus, anterior insula, dorsal anterior cingulate cortex (dACC), dorsolateral prefrontal cortex (dlPFC), and ventromedial prefrontal cortex (vmPFC). Anatomical masks for the amygdala, hippocampus, caudate, putamen, pallidum, ventral striatum, and thalamus were derived from the Harvard–Oxford atlas (Desikan et al., 2006) using a 0.5 probability threshold. Following previous research (Klingelhöfer-Jens, Ehlers, et al., 2022), the anterior insula was defined as the overlap between the thresholded Harvard–Oxford mask (0.5 threshold) and a 60 × 30 × 60 mm box centered at MNI coordinates [0, 30, 0], following anatomical subdivisions (Nieuwenhuys, 2012). Cortical ROIs for the dACC and dlPFC were created by placing a 20 × 16 × 16 mm box around peak voxels identified in a meta-analysis (dACC x-coordinate set to 0; left dlPFC: [−36, 44, 22], right dlPFC: [34, 44, 32], dACC: [0, 18, 42]; Fullana et al., 2016). Following Lonsdorf et al. (2014), the vmPFC mask was constructed as a 20 × 16 × 16 mm box centered on the midline at peak coordinates reported in prior fear learning studies ([0, 40, −12]; e.g., Kalisch et al., 2006; Milad et al., 2007).

#### Data preprocessing

Preprocessing of fMRI data was conducted using SPM12 (Wellcome Department of Neuroimaging, London, UK) running on MATLAB 2013a (The MathWorks, Natick, MA, USA). Preprocessing steps involved discarding the first five volumes of each time series to account for T1 equilibrium effects, slice-timing correction, realignment, coregistration, normalization to a group-specific DARTEL template, and spatial smoothing with a 6 mm full width at half maximum (FWHM) Gaussian kernel.

#### First-level analysis

For the first-level analysis of the acquisition training data, regressors were defined for CS+ and CS-trials. Nuisance regressors included habituation trials, presentation of the experimental screen, US presentations, startle probes, US expectancy ratings and movement parameters. Likewise, for extinction and renewal acquired on day 2, CS+ and CS-were defined as the two regressors of interest. For renewal only the first two trials of each CS were included as trials of interest. The remaining renewal trials, startle probes, US expectancy ratings and movement parameters were again defined as nuisance regressors.

#### Second-level analysis

The first-level CS discrimination contrast (CS+ > CS-) was submitted to second-level one-sample *t*-tests across all experimental phases and time points. Individual anxiety-related trait scores were entered as covariates in the respective models. Whole-brain analyses were conducted using an uncorrected threshold of *p* < .001. In addition, region-of-interest (ROI) analyses were performed for the predefined ROIs. For these analyses, small-volume correction was applied, with significance defined at *p*(FWE) < .05 to account for multiple comparisons.

## Results

### Questionnaires

Internal consistency was excellent with Cronbach’s alpha for STAI-T of 0.92 [95% confidence interval (CI): 0.91, 0.94] and acceptable for NEO-FFI-N with 0.68 [95% confidence interval (CI): 0.63, 0.74] and of 0.75 for IUS [95% confidence interval (CI): 0.71, 0.79]. Distributions of scores are typical for a community sample (see Table 1).

**Table 1.** Pearson correlation coefficients between anxiety-related trait scores and fear-conditioning outcome measures across experimental phases. Values are Pearson correlation coefficients (*r*) with Benjamini–Hochberg (BH)-adjusted *p*-values in parentheses. Significant correlations (*p*BH < .05) are shown in bold. Trend-level correlations (.05 ≤ *p*BH < .10) are shown in italics.

| Measure | Phase | CS discrimination | CS+ | CS- |
| --- | --- | --- | --- | --- |
| SCR (raw) | Acquisition | .128 (.099) | .134 (.099) | .111 (.112) |
|  | Full Extinction | .106 (.135) | .132 (.093) | .147 (.093) |
|  | Early Extinction | .082 (.252) | .130 (.101) | .141 (.101) |
|  | Late Extinction | .103 (.148) | .131 (.096) | .138 (.096) |
|  | Renewal | .091 (.311) | -.016 (.821) | -.145 (.132) |
| SCR (log, rc) | Acquisition | .047 (.505) | .062 (.505) | .054 (.505) |
|  | Full Extinction | .021 (.788) | .058 (.679) | .065 (.679) |
|  | Early Extinction | .009 (.904) | .054 (.725) | .064 (.725) |
|  | Late Extinction | .016 (.836) | .045 (.836) | .053 (.836) |
|  | Renewal | .036 (.693) | .066 (.693) | .034 (.693) |
| SCR (sqrt, z) | Acquisition | .072 (.420) | .057 (.420) | -.078 (.420) |
|  | Full Extinction | -.013 (.877) | -.012 (.877) | .011 (.877) |
|  | Early Extinction | -.012 (.934) | .006 (.934) | .033 (.934) |
|  | Late Extinction | -.018 (.824) | -.044 (.824) | -.016 (.824) |
|  | Renewal | .012 (.864) | -.031 (.864) | -.065 (.864) |
| FPS (raw) | Acquisition | .027 (.926) | .017 (.926) | .007 (.926) |
|  | Full Extinction | .114 (.343) | .078 (.418) | .024 (.735) |
|  | Early Extinction | .099 (.395) | .081 (.395) | .041 (.572) |
|  | Late Extinction | .137 (.172) | .085 (.362) | -.009 (.905) |
|  | Renewal | .148 (.119) | .035 (.628) | -.062 (.590) |
| US expectancy ratings | Acquisition | -.065 (.450) | .002 (.980) | .107 (.255) |
|  | Full Extinction | .031 (.618) | **.145 (.030)** | **.173 (.016)** |
|  | Early Extinction | .018 (.777) | *.123 (.073)* | **.163 (.027)** |
|  | Late Extinction | .050 (.422) | **.153 (.021)** | **.173 (.016)** |
|  | Renewal | .039 (.536) | **.152 (.023)** | **.184 (.010)** |
| Fear ratings | Acquisition | *.131 (.060)* | **.168 (.025)** | .066 (.293) |
|  | Extinction | **.140 (.026)** | **.218 (.001)** | **.170 (.010)** |
|  | Renewal | .049 (.441) | **.204 (.002)** | **.214 (.002)** |

**Note.** SCR = skin conductance response; FPS = fear-potentiated startle; log = log-transformed; rc = range-corrected; sqrt = square-root-transformed; z = z-standardized.

### Main effect of task

Successful acquisition training was evident in all outcome measures (SCR, FPS, US expectancy and post-acquisition fear ratings) with responses to the CS+ being on average significantly higher than to the CS-(all *p*’s < .001; see Figure 4). At a neural level, activation reflecting CS discrimination was observed among others, bilaterally in the insula, the anterior cingulate cortex and supplementary motor area (SMA) as well as the ventral striatum (including the caudate nucleus and pallidum), thalamus and the pons in line with meta-analytic results (Fullana et al., 2016; unthresholded statistical maps are available on NeuroVault: https://identifiers.org/neurovault.collection:24418).

In addition, across all outcome measures (SCR, FPS, US expectancy ratings), responses the CS+ remained significantly greater than responses to the CS-during both early and late extinction (all *p*’s < .001) indicating that differential responding persisted throughout extinction and that extinction was not complete. The decrease in responding from early to late extinction, however, was significantly more pronounced for the CS+ than the CS-(stimulus × time interaction, all *p*’*s* < .001), indicating a reduction in CS discrimination over the course of extinction training. At the neural level, CS discrimination during extinction was associated with activation in the insula, midcingulate cortex, SMA, and caudate nucleus, largely resembling the activation pattern observed during acquisition and consistent with previous meta-analytic findings for extinction (Fullana et al., 2018). In addition, extinction-related CS discrimination was observed in the vmPFC, a region consistently implicated in fear extinction (Gottfried & Dolan, 2004; Milad et al., 2007; Phelps et al., 2004)

During the full renewal phase, responses to the CS+ were larger as compared to the CS-for all outcome measures (SCR, FPS, US expectancy and post-renewal fear ratings; all *p*’s <0.001) (see Figure 2). At the neural level, the renewal effect (CS+ > CS-) in the first two trials was evident only in the insula.

**Figure 2.**
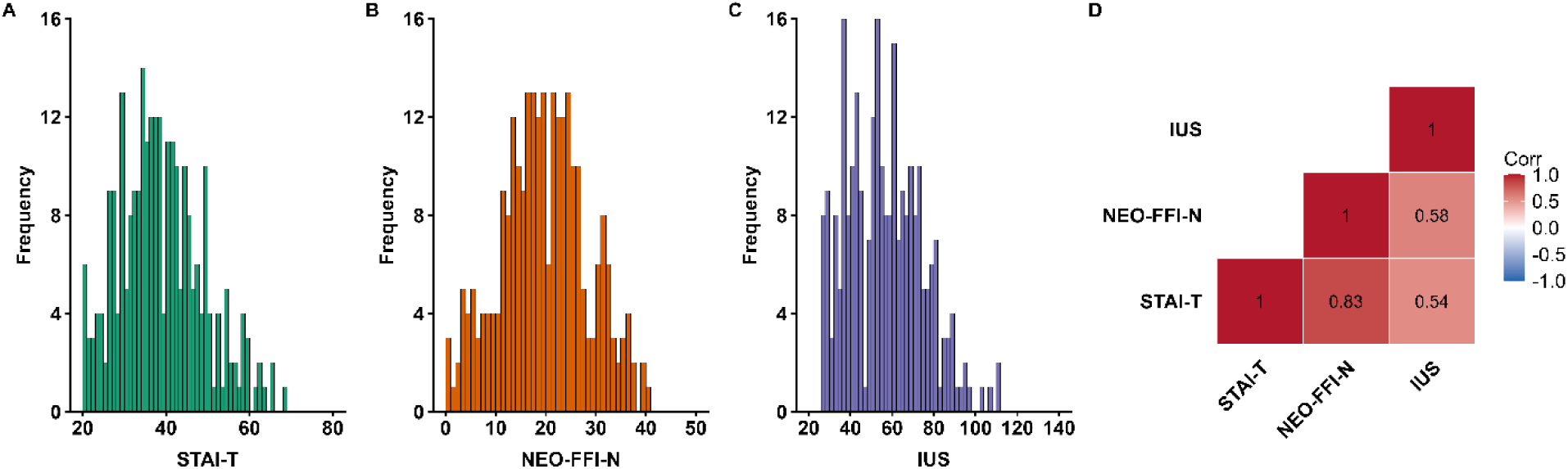
Histograms of individual participant raw scores for STAI-T (A), NEO-FFI-N (B), and IUS (C) and Pearson correlations between STAI-T, NEO-FFI-N, and IUS scores (D).

### Association with anxiety-related traits

Overall, the results indicated no associations between anxiety-related traits and CS discrimination as an index of associative learning in physiological measures (raw or with different transformations of SCR and FPS) or subjective ratings (US expectancy and fear ratings) during any experimental phase (all BH-adjusted *p*’s > .059). Similarly, SCR responses to the US were not associated with ARTs (*r* = .076, *p* = .279).

In contrast, significant positive correlations with responses to both the CS+ and CS-were observed for US expectancy and fear ratings during extinction training and renewal, whereas no corresponding associations were observed for the physiological (SCR) or behavioral measures (FPS) (all BH-adjusted *p*’s > .091). Thus, higher ART scores were associated specifically with higher subjective fear and US expectancy irrespective of CS type, rather than stronger CS discrimination. An overview of correlation coefficients and BH-adjusted *p*-values is provided in Table 1. Corresponding scatterplots are shown in Figure 3. Correlations with individual anxiety-related traits can be found in Supplementary Table 3.

**Figure 3.**
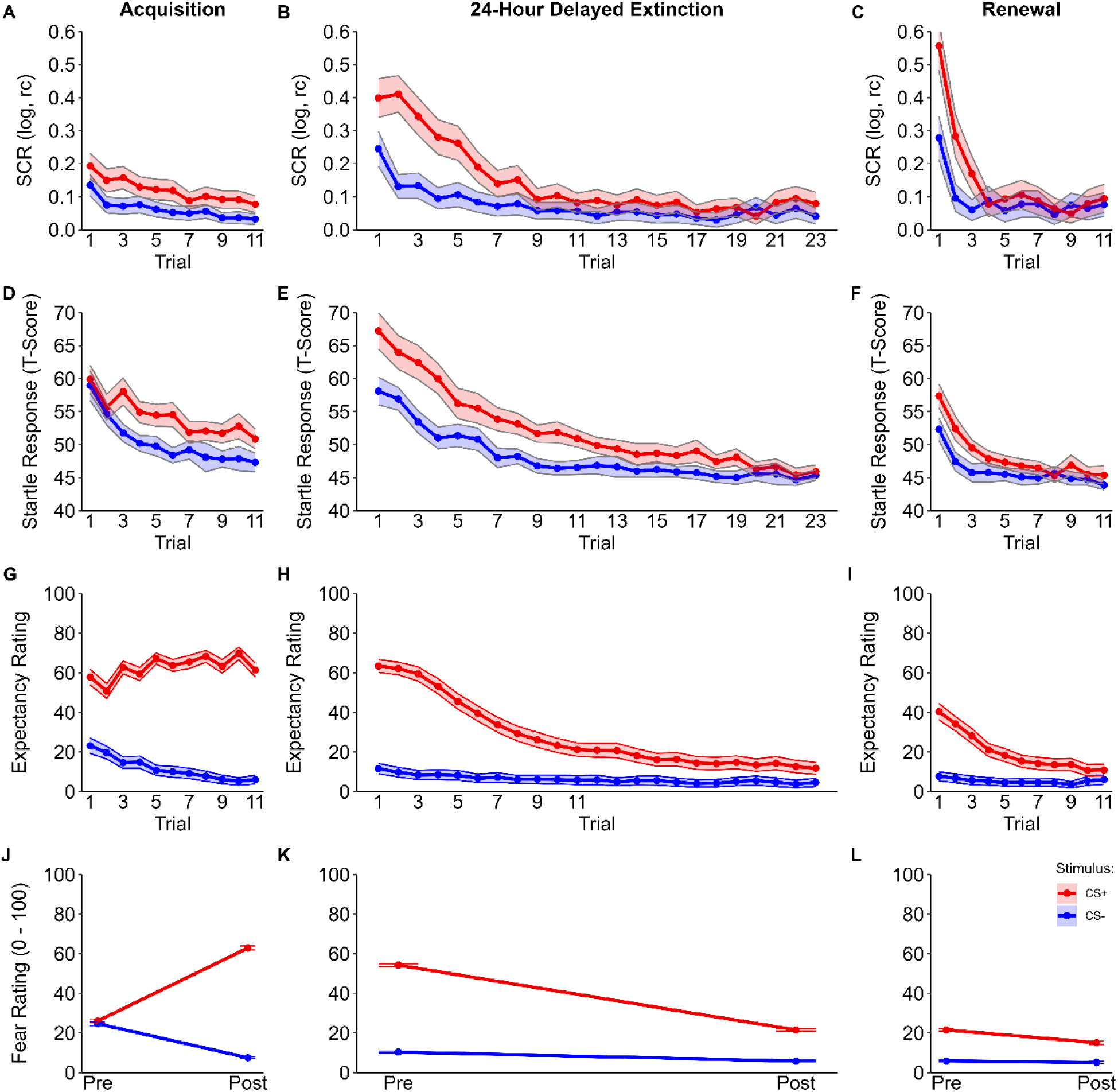
Main effect of task. Trial-by-trial responses during acquisition, extinction, and renewal for (A–C) skin conductance response (SCR), (D–F) fear-potentiated startle (FPS), and (G–I) US expectancy ratings, comparing CS+ and CS-as well as pre and post phase fear ratings (J–L). Shaded colored bands represent 95% confidence intervals.

### ROI analysis

The regions of interest analyses across all phases revealed a significant negative correlation of anxiety-related traits and CS discrimination only in the dorsal anterior cingulate cortex (MNI: −6, 24, 48; t = 3.87, k = 90, *p* = .017 FWE corrected) in the first two trials of renewal as well as nonsignificant trend level negative associations in the right thalamus (MNI: 15, −8, 16; t = 3.76, k = 14, *p* = .060 FWE corrected) in the left anterior insula (MNI: −38, 8, 24; t = 3.57, k = 24, *p* = .074 FWE corrected) for the same contrast (see Figure 4).

**Figure 4.**
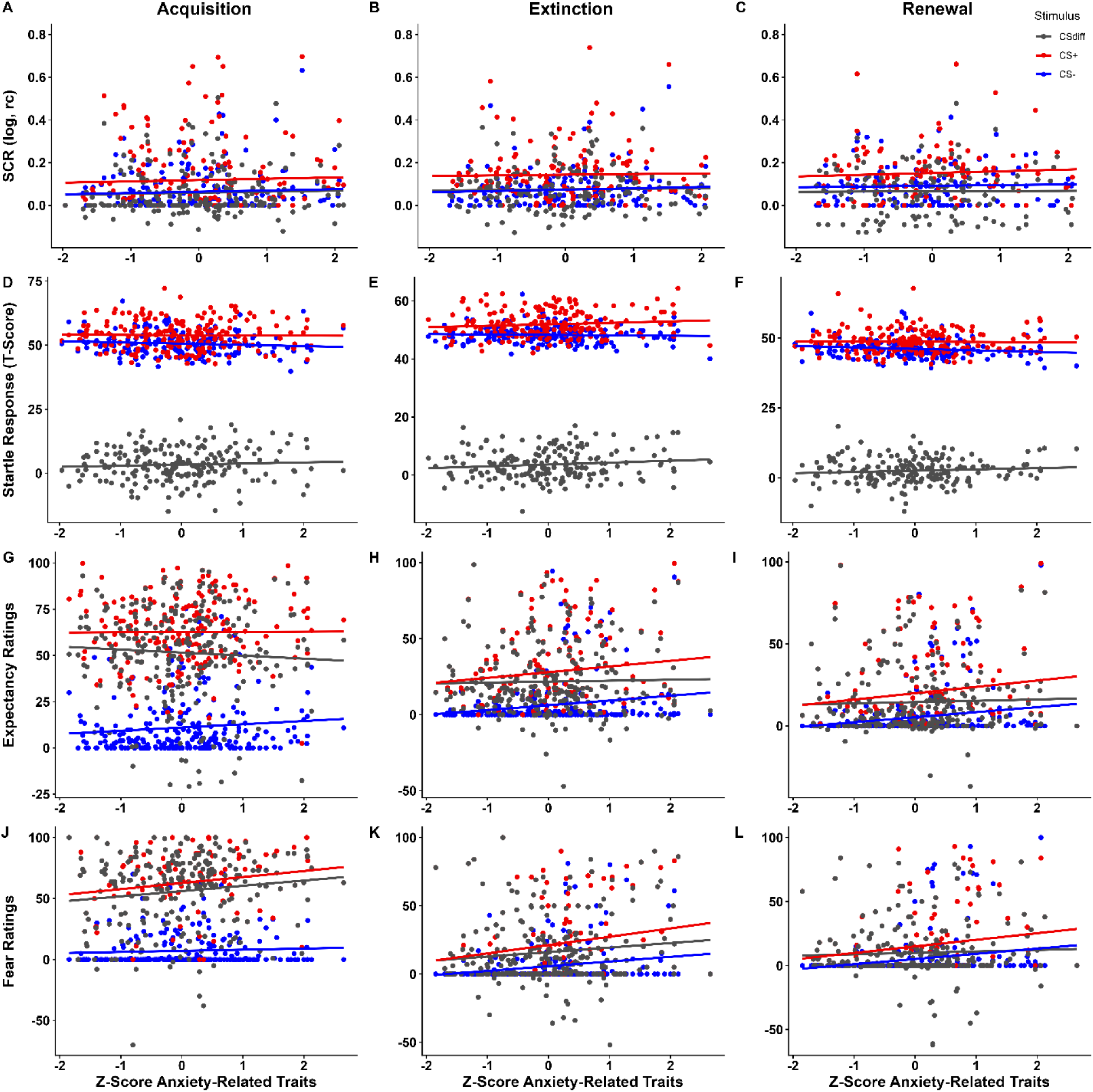
Scatterplots showing associations between anxiety-related traits and responses (CS+ in red, CS-in blue, CS discrimination in grey) during acquisition (A, D, G, I) extinction (B, E, H, K) and renewal (C, F, I, L) for different outcome measures, i.e., SCR (A–C), FPS (D–F), US expectancy ratings (G–I), and fear ratings (J–L).

### Exploratory whole-brain analysis

In an exploratory whole-brain analysis with a voxel-wise threshold of *p* < .001 (uncorrected) and a minimum cluster size of k = 10 voxels, we observed correlations between anxiety-related traits and CS discrimination in several brain regions typically associated with threat processing. During acquisition and late extinction we found a negative correlation in the subgenual part of the anterior cingulate cortex (see Figure 5A, B) and during acquisition and renewal in the orbitofrontal gyrus (see Figure 5C, D). In addition, during late extinction and renewal only, negative correlations could be observed in the anterior insula (see Figure 5E, F) and the thalamus (see Figure 5G, H). These exploratory results imply non-significantly reduced stimulus discrimination in individuals with higher anxiety-related trait scores in brain regions typically linked to successful CS discrimination in previous original and meta-analytical work (for full results see Supplementary Table 4).

**Figure 5.**
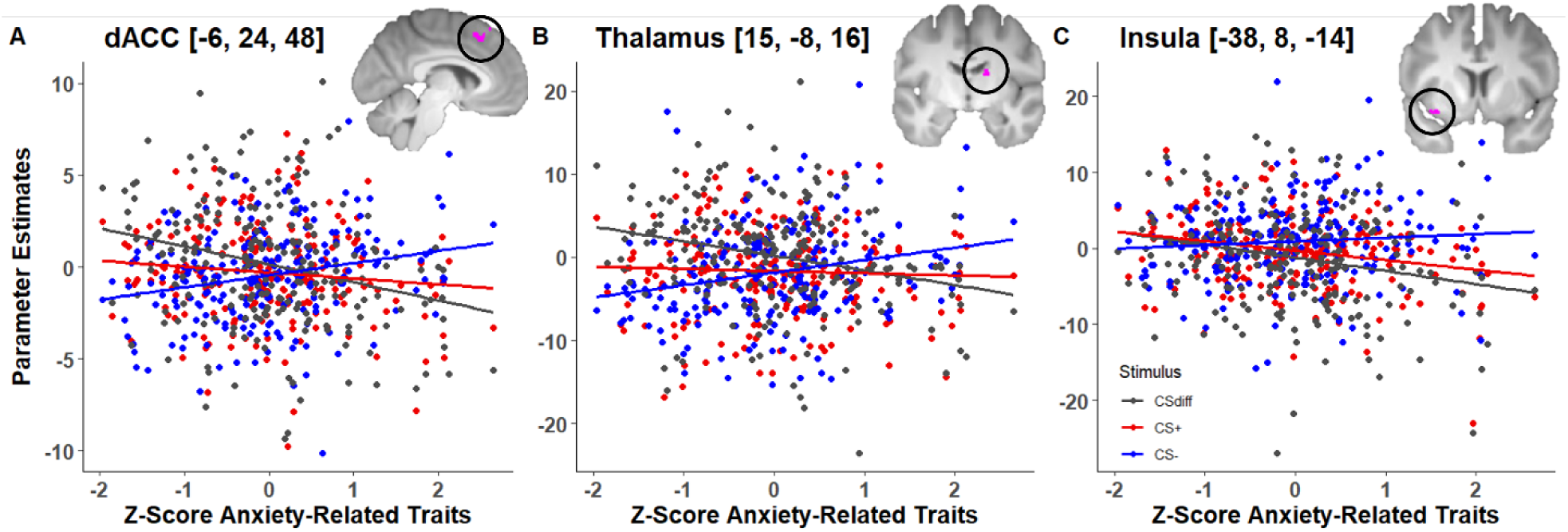
The statistical analysis was based on the CS+ > CS-contrast (CSdiff). Parameter estimates for the CS+ and CS-are additionally shown for illustrative purposes. The visualization threshold was set to *p* = .01, uncorrected.

**Figure 6.**
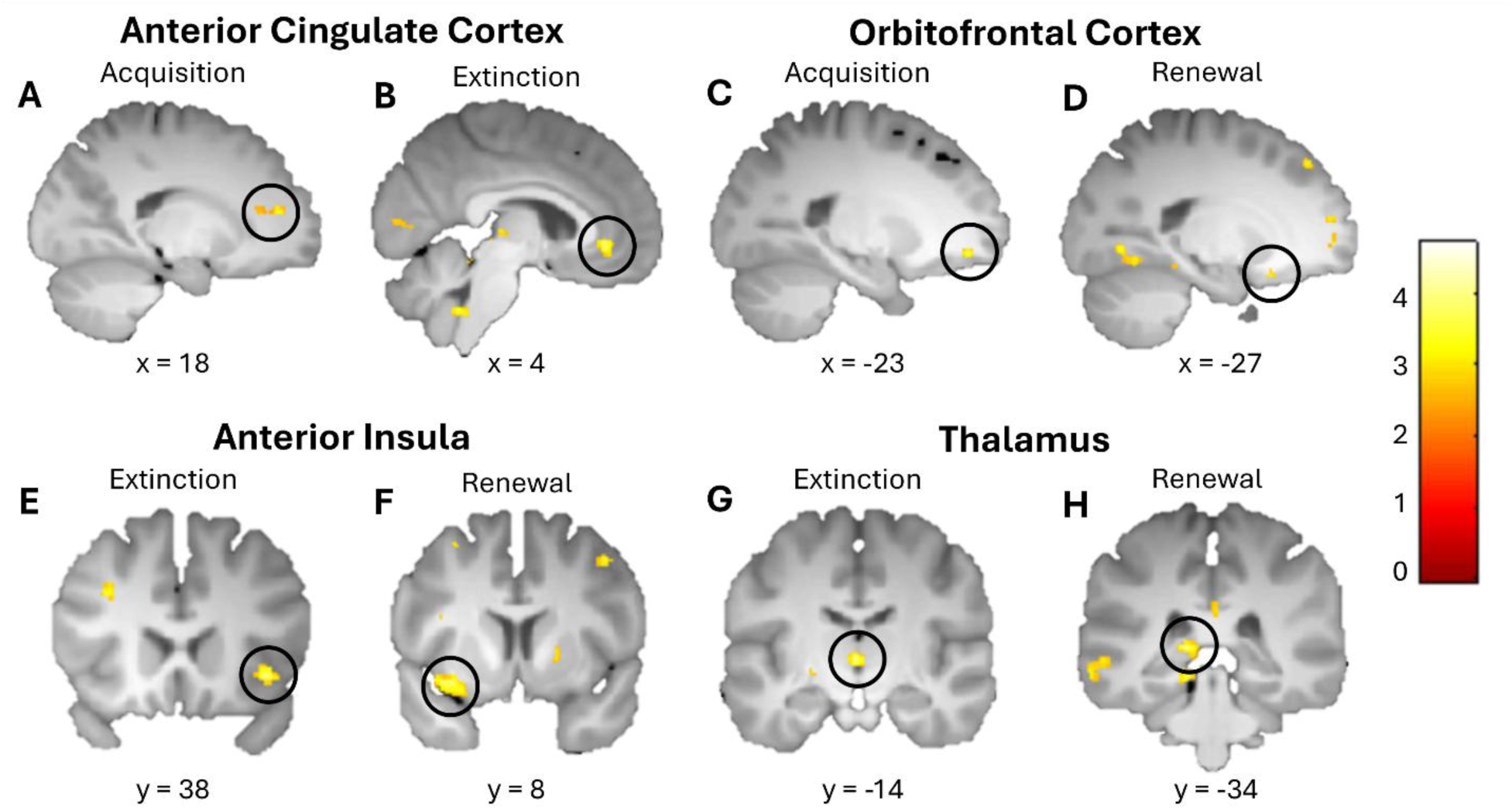
Selected clusters from an exploratory whole brain analysis (*p* < .001 uncorrected) on the negative association between anxiety-related traits and CS+ > CS-contrast. Visualization threshold was set to *p* = .01 uncorrected.

## Discussion

The present study aimed to provide a comprehensive examination of the relationship between anxiety-related traits (ARTs) and associative fear conditioning across acquisition, extinction, and renewal using multimodal outcome measures in a well-powered two-day differential conditioning paradigm. Across experimental phases, ARTs were not associated with CS discrimination as an index of associative learning in either physiological or behavioral measures (SCR and FPS) or subjective ratings (US expectancy and fear ratings) and neural associations were limited, with higher ARTs being related only to significantly reduced CS discrimination in the dACC during early renewal. Importantly, however, higher ARTs were consistently associated with generally (i.e., CS unspecifically) elevated subjective fear and US expectancy ratings during extinction and renewal but not during acquisition training.

Contrary to our hypotheses, across experimental phases, ARTs were not associated with CS discrimination in SCR, FPS, or subjective ratings, providing little evidence for altered associative fear learning with higher ARTs. This pattern largely extended to the neural level, where associations with CS discrimination were limited to reduced dACC discrimination during early renewal, accompanied by trend-level effects in the thalamus and anterior insula, consistent with previous reports of limited or absent neural associations with ARTs (Wendt & Morriss, 2022; Wroblewski et al., 2022). This finding is consistent with several previous studies reporting no associations between ARTs and associated fear learning across phases (e.g., Klingelhöfer-Jens, Morriss, et al., 2022; Mertens & Morriss, 2021; Pineles et al., 2009), but contrasts with reports of reduced CS discrimination during acquisition (e.g., Morriss, Macdonald, et al., 2016; Sjouwerman et al., 2020) or persistent differential responding during extinction (e.g., Morriss, Christakou, et al., 2016; Morriss et al., 2015; Wake et al., 2020), including enhanced neural CS discrimination during extinction (Morriss et al., 2015) – findings that have been interpreted as evidence for deficient safety learning or impaired extinction learning, respectively. Thus, the present findings provide little support for alterations in associative threat learning as a function of ART expression across outcome measures and learning phases. Of note, however, CS-US contingencies were explicitly instructed prior to acquisition training, creating a strong experimental situation that may have constrained interindividual variability and thereby reduced the opportunity for ART-related differences in conditioned responding to emerge during acquisition training (Lissek et al., 2006; Lonsdorf et al., 2017; Mertens et al., 2018) – yet not during extinction training and renewal.

Despite the absence of associations with CS discrimination, higher ARTs were consistently associated with elevated US expectancy and fear ratings to both conditioned stimuli during extinction and renewal. This CS unspecific pattern suggests a cognitive bias in the anticipation and evaluation of threat. Individuals with higher ARTs expected the aversive outcome to occur more strongly irrespective of whether a cue had previously signaled threat or safety and, in turn, reported greater fear of both cues. Related findings indicate that higher trait anxiety is associated with elevated subjective fear not only to the CS+ in an extinction context but also to the CS-across contexts, suggesting that anxiety-related elevations in subjective threat evaluation may extend beyond conditioned threat cues (Haaker et al., 2015). Importantly, the pattern identified in the current study, closely resembles findings in clinical populations. A recent meta-analysis found no differences in CS discrimination between individuals with anxiety-and stress-related disorders and healthy controls, but consistently elevated responses to the individual conditioned stimuli (Kausche et al., 2025). Specifically, during extinction and extinction recall, patients reported heightened threat expectancy and affect ratings to the CS-as well as increased affect ratings to the CS+. Thus, the CS unspecific expectancy and evaluative biases associated with dimensional ARTs in the present non-clinical sample closely parallel the pattern of heightened subjective responding observed in clinical anxiety. This convergence suggests that a heightened tendency to expect and negatively evaluate potential threat, even in the absence of differences in threat–safety discrimination, may represent a process spanning the continuum from anxiety-related traits to anxiety-related psychopathology.

Importantly, this pattern is unlikely to reflect impaired extinction or safety learning in the narrow sense commonly discussed in the anxiety literature (Craske et al., 2022; Duits et al., 2015), as higher ARTs were not associated with altered threat–safety discrimination. Rather, this pattern may further elucidate the cognitive response bias toward threat described above. In another large sample (*N* = 377), Haaker et al. (2015) similarly observed associations between trait anxiety and subjective, but not physiological, responding and suggested that this may reflect a lower threshold for threat appraisal or reduced trust in safety signals. Predictive-processing accounts provide one potential explanation for this pattern: perception is shaped by the integration of prior expectations with current sensory information, with mismatches generating prediction errors that can update these expectations (Van den Bergh et al., 2021). Individuals higher in anxiety may terminate this updating process prematurely and settle on a threat-consistent interpretation, particularly under uncertainty – a “better-safe-than-sorry” strategy that facilitates rapid threat detection but may reduce confidence in safety (de Jong & Vroling, 2014). Rather than reflecting impaired associative learning, higher ARTs may thus be associated with a lower threshold for evaluating potential threat, whereby stimuli are judged as relatively more threatening despite preserved differentiation between threat and safety cues.

The relatively consistent elevation across both conditioned stimuli, together with its restriction to subjective rather than physiological responses, appears most consistent with a broader cognitive or evaluative bias. Other, not mutually exclusive processes may also contribute to this pattern, although explanations commonly discussed in the fear-conditioning literature appear less well suited to account for the complete pattern observed here. Specifically, elevated responding to the CS-could reflect broader generalization of threat appraisal (Cooper et al., 2022), whereas heightened CS+ responding during extinction could be consistent with impaired updating of the CS+–US association, as previously observed in anxiety patients (Rabinak et al., 2017). Elevated responding to both conditioned stimuli may also be compatible with sensitization (Haddad et al., 2012; Naragon-Gainey, 2010) or heightened general threat reactivity (Grillon, 2008), although these processes might be expected to extend beyond subjective responding.

An important strength of the present study is that it simultaneously assessed multiple anxiety-related traits across acquisition training, extinction training, and renewal using physiological, subjective, and neural outcome measures. This comprehensive approach revealed that, while ARTs were not associated with CS discrimination across outcome measures, associations with the general level of responding differed between response systems. Higher ARTs were consistently related to CS unspecific elevations in subjective US expectancy and fear ratings, whereas no comparable general elevations emerged for physiological indices and neural associations were limited. This pattern supports accounts suggesting that different response systems may provide complementary information about conditioned fear rather than interchangeable measures of the same underlying process (Hamm & Vaitl, 1996; Lonsdorf et al., 2017; Ojala & Bach, 2020). Accordingly, individual differences in anxiety-related traits may become more apparent in the general level of responding in outcome measures that rely on explicit evaluations than in autonomic indices of conditioned responding. Consistent with this account, several previous studies have indicated stronger or more robust associations between anxiety-related traits and subjective ratings than with physiological measures (Haaker et al., 2015; Klingelhöfer-Jens, Morriss, et al., 2022; Mertens et al., 2022). A similar pattern was suggested by a meta-analysis of fear generalization (Sep et al., 2019) and has been observed in related blocking and conditioned-inhibition paradigms (Boddez et al., 2012; Kindt & Soeter, 2014). However, as outlined in the Introduction and summarized in Supplementary Table 1, associations across response systems have been heterogeneous, and this pattern has not been observed consistently.

This response-system specificity should not be interpreted as evidence for a strict dissociation between subjective and physiological fear responses. Rather, they support the notion that different outcome measures place different emphasis on partially distinct components of conditioned responding. In this sense, the present results are compatible with perspectives proposing that subjective evaluations and autonomic responses reflect overlapping but not identical processes (Cuve et al., 2023), without necessarily supporting a strict dual-process account of fear learning (LeDoux & Pine, 2016). A further consideration is that anxiety-related traits are themselves predominantly assessed using self-report questionnaires. Their stronger correspondence with subjective evaluative measures than with physiological indices may therefore partly reflect shared variance related to the assessment method. At the same time, subjective experience and physiological responding represent partly distinct components of emotional responding and frequently show limited convergence (Mauss & Robinson, 2009). In summary, these findings underscore the importance of employing multimodal assessment strategies when investigating individual differences in fear conditioning, as conclusions regarding anxiety-related traits may depend on the response system under investigation.

The present findings also highlight the importance of distinguishing between CS discrimination and the general level of responding when characterizing anxiety-related individual differences. A selective focus on CS discrimination may obscure cognitive biases that similarly affect responses to threat and safety cues and therefore cancel out in a difference score, while differences in discrimination alone do not reveal whether they arise from altered responding to threat cues, safety cues or both. At the same time, the manifestation of anxiety-related individual differences may depend on the experimental conditions under which threat learning occurs. Variation in reinforcement rates, contingency instructions, and other procedural characteristics can influence perceived uncertainty and the strength of the experimental situation, potentially enhancing or constraining the expression of trait-related differences (Lissek et al., 2006; Lonsdorf et al., 2017). Thus, variation across previous findings may partly reflect differences in both what aspect of responding was examined and the conditions under which anxiety-related differences had the opportunity to emerge.

Another challenge in integrating findings across studies in the literature concerns differences in how anxiety-related individual differences are conceptualized and measured. Studies have focused on constructs such as trait anxiety, neuroticism, or intolerance of uncertainty, which are conceptually related and share substantial variance but also retain construct-specific components (Sjouwerman et al., 2020). Consequently, findings attributed to different anxiety-related constructs across studies may partly reflect their shared variance, while apparent differences may also arise from construct-specific components. Our exploratory factor analyses support this distinction between shared and construct-specific variance (see Supplementary material for details). STAI-T and NEO-FFI-N items predominantly loaded on a common factor, consistent with the close conceptual and empirical relationship between trait anxiety and neuroticism (Barlow et al., 2014). In contrast, IUS items loaded on a distinct, albeit correlated, factor, consistent with evidence that intolerance of uncertainty is closely related to trait anxiety but captures partly distinct variance (Jensen et al., 2016) and is conceptually more specifically related to negative responses to uncertainty and unpredictability (Carleton, 2016). The additional EFA including measures of anxiety sensitivity and depressive symptoms provided a more nuanced picture of the STAI-T. While most STAI-T items continued to load together with neuroticism, some also loaded on factors capturing depressive symptoms. This pattern is consistent with previous evidence that the STAI-T captures broader negative emotionality, including aspects of neuroticism and depression, rather than anxiety specifically (Bados et al., 2010; Balsamo et al., 2013; Bieling et al., 1998; Knowles & Olatunji, 2020). Thus, the ART measures examined here cannot be regarded as either fully distinct or interchangeable. However, given the large number of items relative to the present sample size, the exploratory factor structure should be interpreted with caution and replicated in a substantially larger sample. Such work will also be important for determining whether associations with CS unspecific expectancy and evaluative biases are driven by shared or construct-specific variance, and ultimately, for identifying the psychological characteristics underlying these biases.

The present findings also raise important questions for future research. In particular, it remains unclear which psychological processes underlie the CS unspecific expectancy and evaluative biases associated with ARTs, to what extent these biases reflect shared versus construct-specific components of anxiety-related traits, and under which conditions they emerge. Addressing these questions is important for determining whether such biases represent clinically relevant processes that contribute to vulnerability to or maintenance of anxiety-related psychopathology. A registered systematic review and meta-analysis currently underway will provide a broader synthesis of associations between anxiety-related traits and fear conditioning and examine methodological variation across studies as well as the content overlap of commonly used anxiety-related questionnaires through item-level analyses (Bruntsch et al., 2024). Ultimately, prospective and clinical studies will be needed to establish whether the cognitive biases identified here predict maladaptive threat responding and the development or persistence of anxiety-related symptoms.

In conclusion, higher anxiety-related traits were not associated with altered CS discrimination across physiological or subjective measures and showed only limited associations with neural indices of conditioned responding. Instead, higher ARTs were consistently associated with CS unspecific elevations in subjective US expectancy and fear during extinction and renewal, suggesting cognitive biases toward heightened threat expectancy and evaluation rather than alterations in associative fear learning. Notably, this pattern closely resembles the heightened subjective responding in the absence of altered CS discrimination observed in individuals with anxiety-and stress-related disorders, suggesting that such biases may represent a process spanning dimensional anxiety-related traits and clinical psychopathology. Together, these findings underscore the importance of distinguishing CS-specific conditioned responding from general levels of subjective responding and of considering multiple response systems when investigating anxiety-related individual differences. Future research should determine the cognitive mechanisms underlying these CS unspecific biases and whether they prospectively predict the development or maintenance of anxiety-related psychopathology.

## CREDIT statement

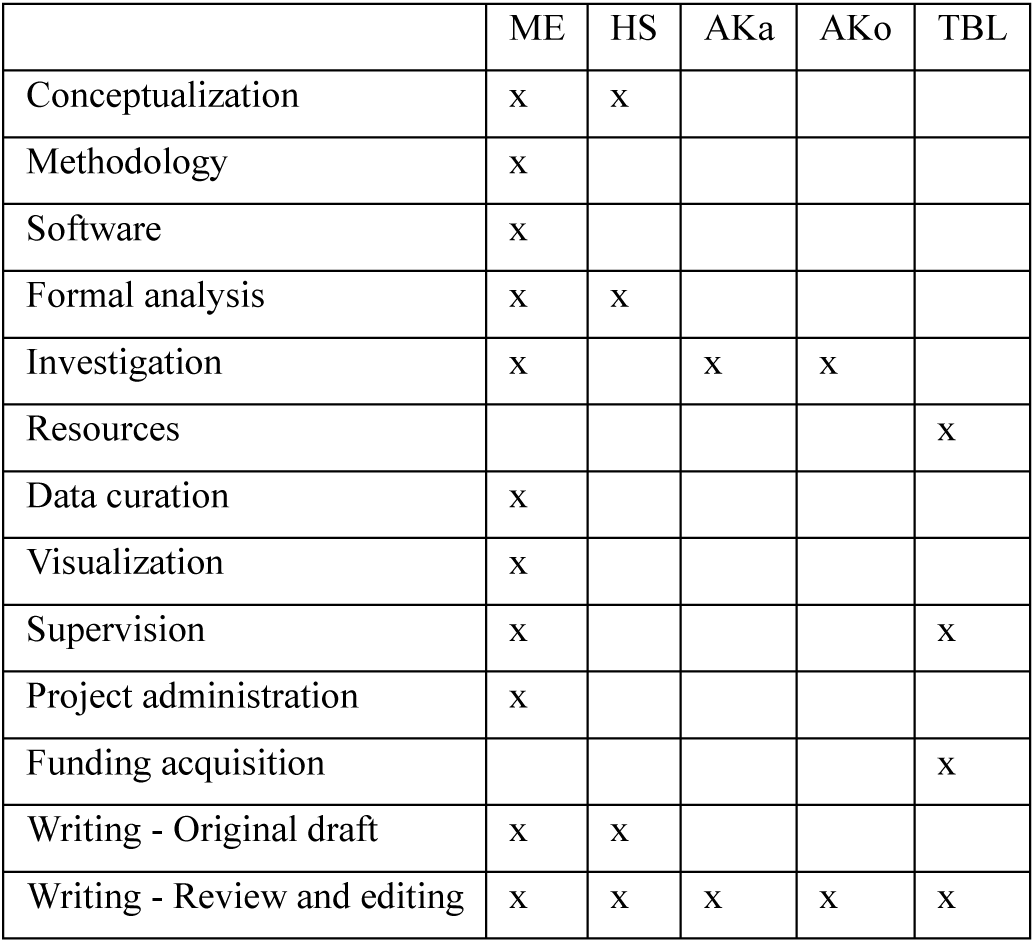

