## Supplementary Material for "Anxiety-Related Traits Are Associated with Subjective Biases but not Altered Threat–Safety Discrimination"

This supplementary section includes additional analyses, figures, and detailed information that support the findings outlined in the main text:

### Anxiety-Related Traits Are Associated with Subjective Biases but not Altered Threat–Safety Discrimination

Mana R. Ehlers, Hannah Stiffel, Alexandros Kastrinogiannis, Alina Koppold, Tina B. Lonsdorf

**Samples sizes**

Two participants did not receive any US presentations during acquisition because of a technical error and were excluded from all fear-conditioning analyses. For SCR, participants were classified as nonresponders and excluded from all SCR analyses if they did not show a measurable response to at least two thirds of the US presentations. For FPS, participants were excluded from all FPS analyses when more than two thirds of their non-missing startle responses during habituation and acquisition were zero. This criterion was evaluated across both CS and intertrial-interval trials. Beyond these measure-specific exclusions, data were occasionally unavailable for individual experimental phases because of technical problems, incomplete recordings, or missing responses. Exclusions were applied only to the affected measure or phase.

| **Measure** | **Acquisition** | **Extinction** | **Renewal** |
|---|---|---|---|
| SCR | 211 | 170 | 139 |
| FPS | 208 | 197 | 196 |
| US-expectancy ratings | 264 | 260 | 258 |
| Fear ratings | 254 | 255 | 256 |
| fMRI | 243 | 222 | 222 |

**Supplementary Table 1.** Overview of included variables and reported results in studies on anxiety-related traits in fear conditioning

+ positive ART-related finding
– negative ART-related finding
n.s. no significant ART-related finding
n/a not reported
no not assessed (inferred as no mentioning in the methods and results)

| | | | | Outcome measure | | | | |
|---|---|---|---|---|---|---|---|---|
| **Author** | **Experimental phase** | **Questionnaires** | **Stimulus** | **SCR** | **FPS** | **US expectancy ratings** | **Fear/distress ratings** | **fMRI** |
| Arnaudova et al. (2017) | Acquisition | EPQ-N[a] | CS+ | n/a | n/a | n/a | no | no |
| | | | CS- | n/a | n/a | n/a | no | no |
| | | | CS discrimination | n.s | n.s | n.s | no | no |
| Chin et al. (2016) | Acquisition | IUS[b] (27-Item-version) | CS+ | no | n/a | no | no | no |
| | | | CS- | no | n/a | no | no | no |
| | | | CS discrimination | no | + | no | no | no |
| Fredrikson & Georgiades (1992) | Acquisition | EPI[c] | CS+ | n/a | no | no | no | no |
| | | | CS- | n/a | no | no | no | no |
| | | | CS discrimination | n.s | no | no | no | no |
| | Extinction | EPI[c] | CS+ | n/a | no | no | no | no |
| | | | CS- | n/a | no | no | no | no |
| | | | CS discrimination | n.s | no | no | no | no |
| Gazendam et al. (2013) | Acquisition | STAI-T[d] | CS+ | n.s | n.s | n.s | n.s | no |
| | | | CS- | n.s | + | n.s | + | no |
| | | | CS discrimination | n.s | –[1] | – | –[1] | no |
| | Extinction | STAI-T[d] | CS+ | n.s | n.s | n.s | + | no |
| | | | CS- | n.s | n.s | n.s | + | no |
| | | | CS discrimination | n.s | n.s | n.s | n.s | no |
| Haaker et al. (2015) | Acquisition | STAI-T[e] | CS+ | n.s | no | n.s | n.s | no |
| | | | CS- | n.s | no | n.s | + | no |
| | | | CS discrimination | n/a | no | n/a | n/a | no |
| | Extinction | STAI-T[e] | CS+ | n.s | no | + | + | no |
| | | | CS- | n.s | no | n.s | + | no |
| | | | CS discrimination | n/a | no | n/a | n/a | no |
| Haddad et al. (2012) | Acquisition | STAI-T[e] | CS+ | n.s | n.s | no | + | no |
| | | | CS- | n.s | n.s | no | + | no |
| | | | CS discrimination | n.s | – | no | n.s | no |
| | Extinction | STAI-T[e] | CS+ | n.s | n.s | no | n.s | no |
| | | | CS- | n.s | n.s | no | n.s | no |
| | | | CS discrimination | n.s | n.s | no | n.s | no |

| | | | | Outcome measure | | | | |
|---|---|---|---|---|---|---|---|---|
| **Author** | **Phase** | **Questionnaires** | **Stimulus** | **SCR** | **FPS** | **US expectancy ratings** | **Fear/distress ratings** | **fMRI** |
| Hur et al. (2016) | Acquisition | GTS[f] | CS+ | n/a | no | no | no | no |
| | | | CS- | n/a | no | no | no | no |
| | | | CS discrimination | n.s | no | no | no | no |
| | Extinction | GTS[f] | CS+ | n/a | no | no | no | no |
| | | | CS- | n/a | no | no | no | no |
| | | | CS discrimination | n.s | no | no | no | no |
| Indovina et al. (2011) | Acquisition | STAI-T[e] | CS+ | n/a | no | no | no | n/a |
| | | | CS- | n/a | no | no | no | n/a |
| | | | CS discrimination | + | no | no | no | + |
| Klingelhöfer-Jens et al. (2022) | Acquisition | STAI-T[e] | CS+ | n.s | n.s | no | + | no |
| | | | CS- | n.s | n.s | no | + | no |
| | | | CS discrimination | n.s | n.s | no | n.s | no |
| | | IUS[b] (27-Item-version) | CS+ | n.s | n.s | no | + | no |
| | | | CS- | n.s | n.s | no | n.s | no |
| | | | CS discrimination | n.s | n.s | no | n.s | no |
| | Extinction | STAI-T[e] | CS+ | n.s | n.s | no | n.s | no |
| | | | CS- | n.s | n.s | no | n.s | no |
| | | | CS discrimination | n.s | n.s | no | n.s | no |
| | | IUS[b] (27-Item-version) | CS+ | n.s | n.s | no | n.s | no |
| | | | CS- | n.s | n.s | no | n.s | no |
| | | | CS discrimination | n.s | n.s | no | n.s | no |
| Lommen et al. (2010) | Acquisition | EPQ-N[a] | CS+ | no | no | n/a | no | no |
| | | | CS- | no | no | n/a | no | no |
| | | | CS discrimination | no | no | n.s | no | no |
| Martínez et al. (2012) | Acquisition | STAI-T[d] | CS+ | n.s | no | no | no | no |
| | | | CS- | n.s | no | no | no | no |
| | | | CS discrimination | n.s | no | no | no | no |
| | | NEO-FFI-N[g] | CS+ | n.s | no | no | no | no |
| | | | CS- | n.s | no | no | no | no |
| | | | CS discrimination | n.s | no | no | no | no |
| | Extinction | STAI-T[d] | CS+ | n.s | no | no | no | no |
| | | | CS- | n.s | no | no | no | no |
| | | | CS discrimination | n.s | no | no | no | no |
| | | NEO-FFI-N[g] | CS+ | n.s | no | no | no | no |
| | | | CS- | n.s | no | no | no | no |
| | | | CS discrimination | n.s | no | no | no | no |
| | Renewal | STAI-T[d] | CS+ | n.s | no | no | no | no |

| | | | | | | | | |
|---|---|---|---|---|---|---|---|---|
| | | | CS- | n.s | no | no | no | no |
| | | | CS discrimination | n.s | no | no | no | no |
| | | NEO-FFI-N[g] | CS+ | n.s | no | no | no | no |
| | | | CS- | n.s | no | no | no | no |
| | | | CS discrimination | n.s | no | no | no | no |

| | | | | Outcome measure | | | | |
|---|---|---|---|---|---|---|---|---|
| **Author** | **Phase** | **Questionnaires** | **Stimulus** | **SCR** | **FPS** | **US expectancy ratings** | **Fear/distress ratings** | **fMRI** |
| Mertens et al. (2022) | Acquisition | IUS[b] (27-Item-version) | CS+ | n.s | n.s | n.s | + | no |
| | | | CS- | n.s | – | n.s | n.s | no |
| | | | CS discrimination | n.s | n.s | n.s | n.s | no |
| Mertens & Morriss (2021) | Acquisition | IUS[h] (12-Item-version) | CS+ | n/a | n/a | no | no | no |
| | | | CS- | n/a | n/a | no | no | no |
| | | | CS discrimination | n.s | n.s | no | no | no |
| Morriss (2019) | Acquisition | STAI-T[e] | CS+ | n/a | no | n/a | no | no |
| | | | CS- | n/a | no | n/a | no | no |
| | | | CS discrimination | n.s | no | n.s | no | no |
| | | IUS[b] (27-Item-version) | CS+ | n/a | no | n/a | no | no |
| | | | CS- | n/a | no | n/a | no | no |
| | | | CS discrimination | n.s | no | n.s | no | no |
| | Late Extinction | STAI-T[e] | CS+ | n/a | no | n/a | no | no |
| | | | CS- | n/a | no | n/a | no | no |
| | | | CS discrimination | n.s | no | n.s | no | no |
| | | IUS[b] (27-Item-version) | CS+ | n/a | no | n/a | no | no |
| | | | CS- | n/a | no | n/a | no | no |
| | | | CS discrimination | +[2] | no | n.s | no | no |
| Morriss et al. (2015) | Acquisition | IUS[h] | CS+ | n/a | no | no | no | n/a |
| | | | CS- | n/a | no | no | no | n/a |
| | | | CS discrimination | n.s | no | no | no | n.s |
| | Late Extinction | IUS[h] | CS+ | n.s | no | no | no | n/a |
| | | | CS- | n.s | no | no | no | n/a |
| | | | CS discrimination | +[2] | no | no | no | + |
| Morriss, Christakou, et al. (2016) | Acquisition | IUS[h] (27-Item-version) | CS+ | n/a | no | no | no | no |
| | | | CS- | n/a | no | no | no | no |
| | | | CS discrimination | n.s | no | no | no | no |
| | Late Extinction | IUS[h] (27-Item-version) | CS+ | n/a | no | no | no | no |
| | | | CS- | n/a | no | no | no | no |
| | | | CS discrimination | +[2] | no | no | no | no |
| Morriss, Macdonald, et al. (2016) | Acquisition | IUS[h] | CS+ | n/a | no | no | no | no |
| | | | CS- | n/a | no | no | no | no |

| | | | | | | | | |
|---|---|---|---|---|---|---|---|---|
| | | | CS discrimination | –[2] | no | no | no | no |
| | Extinction | IUS[h] | CS+ | n/a | no | no | no | no |
| | | | CS- | n/a | no | no | no | no |
| | | | CS discrimination | +[2] | no | no | no | no |

| | | | | Outcome measure | | | | |
|---|---|---|---|---|---|---|---|---|
| **Author** | **Phase** | **Questionnaires** | **Stimulus** | **SCR** | **FPS** | **US expectancy ratings** | **Fear/distress ratings** | **fMRI** |
| Morriss, Saldarini, Chapman, et al. (2019) | Acquisition | IUS[h] | CS+ | n/a | no | n/a | no | no |
| | | | CS- | n/a | no | n/a | no | no |
| | | | CS discrimination | n.s | no | n.s | no | no |
| Morriss et al. (2019), study 1 | Acquisition | STAI-T[e] | CS+ | n/a | no | n/a | no | no |
| | | | CS- | n/a | no | n/a | no | no |
| | | | CS discrimination | + | no | – | no | no |
| | | IUS[b] (27-Item-version) | CS+ | n/a | no | n/a | no | no |
| | | | CS- | n/a | no | n/a | no | no |
| | | | CS discrimination | n.s | no | n.s | no | no |
| | Extinction | STAI-T[e] | CS+ | n/a | no | n/a | no | no |
| | | | CS- | n/a | no | n/a | no | no |
| | | | CS discrimination | n.s | no | – | no | no |
| | | IUS[b] (27-Item-version) | CS+ | n/a | no | n/a | no | no |
| | | | CS- | n/a | no | n/a | no | no |
| | | | CS discrimination | + | no | – | no | no |
| Morriss & van Reekum (2019)[3], study 1 | Acquisition | IUS[h] (27-Item-version) | CS+ | n/a | no | n/a | no | no |
| | | | CS- | n/a | no | n/a | no | no |
| | | | CS+ discrimination | n.s | no | n.s | no | no |
| | Extinction | IUS[h] (27-Item-version) | CS+ | + | no | n.s | no | no |
| | | | CS- | – | no | n.s | no | no |
| | | | CS discrimination | + | no | n.s | no | no |
| Morriss et al. (2020) | Acquisition | STICSA[i] | CS+ | n/a | no | n/a | no | no |
| | | | CS- | n/a | no | n/a | no | no |
| | | | CS discrimination | n.s | no | n.s | no | no |
| | | IUS[b] (27-Item-version) | CS+ | n/a | no | n/a | no | no |
| | | | CS- | n/a | no | n/a | no | no |
| | | | CS discrimination | n.s | no | n.s | no | no |
| | Extinction | STICSA[i] | CS+ | n/a | no | n/a | no | no |
| | | | CS- | n/a | no | n/a | no | no |
| | | | CS discrimination | n.s | no | n.s | no | no |
| | | IUS[b] (27-Item-version) | CS+ | n/a | no | n/a | no | no |
| | | | CS- | n/a | no | n/a | no | no |
| | | | CS discrimination | n.s | no | n.s | no | no |

| Author | Phase | Questionnaires | Stimulus | Outcome measure | | | | |
|---|---|---|---|---|---|---|---|---|
| | | | | **SCR** | **FPS** | **US expectancy ratings** | **Fear/distress ratings** | **fMRI** |
| Otto et al. (2007) | Acquisition | NEO-FFI-N[g] | CS+ | n.s | no | no | no | no |
| | | | CS- | n/a | no | no | no | no |
| | | | CS discrimination | n.s | no | no | no | no |
| Pineles et al. (2009) | Acquisition | NEO-PI-R[j] | CS+ | n.s | no | no | no | no |
| | | | CS- | n/a | no | no | no | no |
| | | | CS discrimination | n.s | no | no | no | no |
| | Extinction | NEO-PI-R[j] | CS+ | n.s | no | no | no | no |
| | | | CS- | n/a | no | no | no | no |
| | | | CS discrimination | n.s | no | no | no | no |
| Sjouwerman et al. (2020), study 1 | Acquisition | STAI-T[e] | CS+ | n.s | n.s | no | n.s | no |
| | | | CS- | n.s | + | no | n.s | no |
| | | | CS discrimination | – | n.s | no | n.s | no |
| | | IUS[h] (27-Item-version) | CS+ | n.s | n.s | no | n.s | no |
| | | | CS- | n.s | + | no | n.s | no |
| | | | CS discrimination | n.s | – | no | n.s | no |
| | | NEO-FFI-N[k] | CS+ | n.s | n.s | no | n.s | no |
| | | | CS- | n.s | + | no | n.s | no |
| | | | CS discrimination | n.s | n.s | no | n.s | no |
| study 2 | Acquisition | STAI-T[e] | CS+ | n.s | no | no | n.s | + |
| | | | CS- | n.s | no | no | + | n.s |
| | | | CS discrimination | n.s | no | no | n.s | + |
| Tzschoppe et al. (2014) | Acquisition | NEO-FFI-N[l] | CS+ | n.s | no | no | n.s | n.a |
| | | | CS- | n.s | no | no | n.s | n.a |
| | | | CS discrimination | n.s | no | no | n.s | +[4] |
| | Extinction | NEO-FFI-N[l] | CS+ | n.s | no | no | n.s | n.a |
| | | | CS- | n.s | no | no | n.s | n.a |
| | | | CS discrimination | n.s | no | no | n.s | n.a |
| Wake et al. (2020) | Acquisition | STICSA[i] | CS+ | n/a | no | n/a | no | no |
| | | | CS- | n/a | no | n/a | no | no |
| | | | CS discrimination | n.s | no | n.s | no | no |
| | | IUS[b] (27-Item-version) | CS+ | n/a | no | n/a | no | no |
| | | | CS- | n/a | no | n/a | no | no |
| | | | CS discrimination | n.s | no | n.s | no | no |
| | Extinction | STICSA[i] | CS+ | n/a | no | n/a | no | no |
| | | | CS- | n/a | no | n/a | no | no |
| | | | CS discrimination | + | no | + | no | no |
| | | IUS[b] (27-Item-version) | CS+ | n/a | no | n/a | no | no |
| | | | CS- | n/a | no | n/a | no | no |

| | | | | | | | | |
|---|---|---|---|---|---|---|---|---|
| | | | CS discrimination | + | no | n.s | no | no |

| | | | | Outcome measure | | | | |
|---|---|---|---|---|---|---|---|---|
| **Author** | **Phase** | **Questionnaires** | **Stimulus** | **SCR** | **FPS** | **US expectancy ratings** | **Fear/distress ratings** | **fMRI** |
| Wendt & Morriss (2022) | Acquisition | IUS[b] (27-Item-version) | CS+ | n/a | no | no | no | n/a |
| | | | CS- | n/a | no | no | no | n/a |
| | | | CS discrimination | n.s | no | no | no | n.s |
| | Extinction | IUS[b] (27-Item-version) | CS+ | n/a | no | no | no | n/a |
| | | | CS- | n/a | no | no | no | n/a |
| | | | CS discrimination | n.s | no | no | no | n.s |
| Wroblewski et al. (2022) | Acquisition | STAI-T[e] & IUS[h] (27-Item-version) | CS+ | + | n.s | n.s | no | no |
| | | | CS- | n.s | n.s | n.s | no | no |
| | | | CS discrimination | n.s | n.s | n.s | no | no |
| | Early Extinction | STAI-T[e] & IUS[h] (27-Item-version) | CS+ | n.s | n.s | n.s | no | – |
| | | | CS- | n.s | n.s | n.s | no | – |
| | | | CS discrimination | + | + | n.s | no | n.s |
| Ehlers et al. | Acquisition | STAI-T[d], NEO-FFI-N[k] IUS[h] (27-Item-version) | CS+ | n.s | n.s | n.s | + | |
| | | | CS- | n.s | n.s | n.s | n.s | |
| | | | CS discrimination | n.s | n.s | n.s | n.s | n.s |
| | Extinction | STAI-T[d], NEO-FFI-N[k] IUS[h] (27-Item-version) | CS+ | n.s | n.s | + | + | |
| | | | CS- | n.s | n.s | + | + | |
| | | | CS discrimination | n.s | n.s | n.s | n.s | n.s |
| | Renewal | STAI-T[d], NEO-FFI-N[k] IUS[h] (27-Item-version) | CS+ | n.s | n.s | + | + | |
| | | | CS- | n.s | n.s | + | + | |
| | | | CS discrimination | n.s | n.s | n.s | n.s | – |

**Note.** The publications listed here do not constitute an exhaustive list of all studies on the research topic of interest. For a comprehensive overview on related studies, see pre-registered meta-analysis in preparation (Bruntsch et al., 2024). Detailed information on design specifications of the listed publications is provided in Supplementary Table 2. Note that trend-level effects are marked as non-significant.

[1] Categorial analysis: The groups with high and low trait scores differed in the course of acquisition. CS-type x Trial x Group interaction, i.e., the acquisition effect, was significant in the control group but absent in the high trait anxiety group.
[2] Different results were reported for groups with high and low trait scores (e.g., CS discrimination was significant for the group with high STAIT scores but not significant for the group with low scores). However, the groups were not tested against each other.
[3] The results are only valid for the uninstructed group in this study.
[4] In this study, higher neuroticism scores have been linked to stronger *functional connectivity* between the amygdala and hippocampus as well as between the amygdala and prefrontal regions during acquisition training.

[a] Eysenck Personality Questionnaire (EPQ-N; Eysenck & Eysenck, 1975)
[b] Intolerance of uncertainty scale (IUS; Freeston et al., 1994)

[c] Eysenck Personality Inventory (EPI; (Eysenck & Eysenck, 1968)
[d] State-Trait Anxiety Inventory (STAI-T; Spielberger et al., 1970)
[e] State-Trait Anxiety Inventory (STAI-T; Spielberger et al., 1983)
[f] Negative Temperament subscale of the General Temperament Survey (GTS; Watson & Clark, 1993)
[g] NEO-Five Factor Inventory (NEO-FFI; (Costa Jr. & McCrae, 1992a)
[h] Intolerance of uncertainty scale (IUS; (Buhr & Dugas, 2002)
[I] State-Trait Inventory for Cognitive and Somatic Anxiety (STICSA; Ree et al., 2008)
[j] Revised NEO Personality Inventory (NEO-PI-R; (Costa Jr. & McCrae, 1992b)
[k] NEO Five-Factor Inventory (NEO-FFI-N; McCrae & Costa, 2004)
[l] NEO Five-Factor Inventory (NEO-FFI-N; Costa Jr. & McCrae, 1997)

**Supplementary Table 2.** Overview on design specifications of studies on anxiety-related traits in fear conditioning.

| Author | N (prior to exclusion) | Conditioning type | Experimental timing | Reinforcement ratio | CS type | Number CS+/ number CS- | US type | Instruction type (CS-US contingency) | Trials acquisition | Trials extinction | ROF type |
|---|---|---|---|---|---|---|---|---|---|---|---|
| Arnaudova et al. (2017) | N=58 | Differential | Single day | 100% | Circles on a black-white continuum | 2/2 | Electrotactile | (c) | 10 (5 each)/ 10 (5 each) | – | – |
| Chin et al. (2016) | N=32 | Differential | Single day | 50% and 75% in 2 blocks | Geometric shapes | 1/1 | Electrotactile | (d) | 16/16 | – | – |
| Fredrikson & Georgiades (1992) | N=46 | Differential | Single day | 100% | Geometric shapes | 1/1 | Electrotactile | (b) | 8/8 | 8/8 | – |
| Gazendam et al. (2013) | N=42 | Differential | Day 1: Acquisition Day2: Extinction, Reinstatement test + Re-extinction | 87.5% | Two male faces with neutral expressions | 1/1 | Electrotactile | Acquisition: (d) Extinction, reinstatement test, re-extinction: (a) | 8/8 | 10/10 | Reinstatement test + Re-extinction |
| Haaker et al. (2015) | N=377 | Differential cue and context | Day 1: Acquisition Extinction (Day 2: Fear and Extinction Recall) | 50% | Geometric shapes | 1/1 | Electrotactile | (a) | 24/24 | 24/24 | – |
| Haddad et al. (2012) | N=50 | Differential | Single day | 75% | Female faces with neutral expression; grey oval as dissimilar CS- | 1/1 | Fearful face and scream (95dB) | n/a | 12/12/12 | 9/9/9 | – |
| Hur et al. (2016) | N=97 | Differential | Single day | 100% | Fear relevant stimuli (spider, snake), non-fearful control stimuli (mushrooms) | 2/2/5 (control) | Electrotactile | n/a | 12/12/15 | 16/16/20 | – |
| Indovina et al. (2011) | N=23 | Combined context-cue conditioning | Day 1: Training/Acquisition Day 3:fMRI | Predictable CS: 100% | Virtual actor (m or f) putting their hands to their ears | CS predictable, non-predictable, safe | Scream (103dB) | n/a | Day 1 (behavioral lab): 5 each room Day 3 (fMRI): 16 each room | – | – |
| Klingelhöfer-Jens et al. (2022) | N=66 | Differential | Day 1: Acquisition Day 2: Extinction, Reinstatement test | 100% | White snow fractals | 1/1 | Electrotactile | (a) | 9/9 | 9/9 | Reinstatement test |
| Lommen et al. (2010) | N=55 | Differential | Single day | 100% | 10 circles in degraded grey color | 2/2 | Electrotactile | n/a | 10 (5 each)/ 10 (5 each) | 10 (5 each)/10 (5 each) | –[1] |

| Author | N (prior to exclusion) | Conditioning type | Experimental timing | Reinforcement ratio | CS type | Number CS+/ number CS- | US type | Instruction type (CS-US contingency) | Trials acquisition | Trials extinction | ROF type |
|---|---|---|---|---|---|---|---|---|---|---|---|
| Martínez et al. (2012) | N=46 | Combined context-cue conditioning | Day 1: Acquisition Extinction Day 2: Recall Renewal | 100% | Red and blue desk light (embedded in contexts) | 1/1 | Electrotactile | (b) | 5/5 | 10/10 | Recall, Renewal |
| Mertens et al. (2022) | N=120 | Differential | Single day | 75% | Five words related to size | 1/1 | Electrotactile | (c) | 8/8 | – | – |
| Mertens & Morriss (2021) | N=108 | Differential | Single day | Acquisition: 75% Reversal: 20% | Geometric shapes | 1/1 | Electrotactile | Acquisition: (b) vs (d) vs (e) Reversal: (e) | 8/8 | – | (Reversal) |
| Morriss (2019) | N=45 | Differential | Single day | 50% | Geometric shapes | 1/1 | Female scream (90dB) | (a) | 12/12 | 16/16 | – |
| Morriss et al. (2015) | N=22 | Differential | Single day | Acquisition: 100% | Geometric shapes | 1/1 | Female scream (90dB) | (a) | 12/12 | 16/16 | – |
| Morriss, Christakou, et al. (2016) | N=38 | Differential | Single day | Acquisition: 100% | Geometric shapes | 1/1 | Female scream (90dB) | (a) | 12/12 | 16/16 | – |
| Morriss, Macdonald, et al. (2016) | N=54 | 1CS+ and 3 CS- (GS1, GS2, GS3) | Single day | 50% | Geometric shapes | 1/3 (generalization CS) | Female scream (90dB) | (b) | 12/8/ GS: 8/8 | 10/10/10/ 10 | – |
| Morriss, Saldarini, Chapman, et al. (2019) | N=44 | Differential | Single day | 50% in acquisition and reversal | Geometric shapes | 1/1 | Female scream (90dB) | (a) | 12/12 | – | (Reversal) |
| Morriss et al. (2019), study 1 | N=30 | Differential | Single day | 50% | Geometric shapes | 1/1 | Female scream (90dB) | (a) | 12/12 | 16/16 | – |
| Morriss & van Reekum (2019)[3], study 1 | N=60 | Differential | Single day | 50% | Geometric shapes | 1/1 | Female scream (90dB) | Acquisition: (a) Extinction: (a) vs (e) | 12/12 | 16/16 | – |
| Morriss et al. (2020) | N=144 | Differential | Single day vs. Day 1: Acquisition Day 2: Extinction | 50% | Geometric shapes | 1/1 | Female scream (90dB) | (a) | 12/12 | Regular: 16/16 Extended: 24/24 | – |
| Otto et al. (2007) | N= 73 | Differential | Single day | 100% | Geometric shapes | 1/1 | Electrotactile | n/a | 5/5 | – | – |
| Pineles et al. (2009) | N= 217 | Differential | Single day | 100% | Geometric shapes | 1/1 | Electrotactile | (b) | 5/5 | 10/10 | – |
| Sjouwerman et al. (2020), study 1 | N=116 | Differential | Single day | 100% | Geometric shapes | 1/1 | Electrotactile | (a) | 9/9 | – | – |
| study 2 | N=124 | Differential | Single day | 100% | White fractals | 1/1 | Electrotactile | (a) | 14/14 | – | – |
| Tzschoppe et al. (2014) | N=47 | Differential | Single day | 50% | Male faces with neutral expression | 1/1 | Female scream (80dB) | (a) | 30/30 | 10/10 | – |
| Wake et al. (2020) | N=95 | Differential | Single day | 50% | Geometric shapes | 1/1 | Female scream (90dB) | Acquisition: (a) Extinction: (a) vs (c) | 12/12 | 16/16 | – |

| Author | N (prior to exclusion) | Conditioning type | Experimental timing | Reinforcement ratio | CS type | Number CS+/ number CS- | US type | Instruction type (CS-US contingency) | Trials acquisition | Trials extinction | ROF type |
|---|---|---|---|---|---|---|---|---|---|---|---|
| Wendt & Morriss (2022) | N=48 | Differential | Single day | 50% | Geometric shapes | 1/1 | Electrotactile | Acquisition: (c) Mid-Acquisition: (e) Mid-Extinction: (b) | 16/16 | 16/16 | – |
| Wroblewski et al. (2022) | N=155 | Differential | Day 1: Acquisition Day 2: Extinction Reinstatement test Re-extinction | 60% | Male faces with neutral expression | 1/1 | Electrotactile | Acquisition: (e) Extinction, Reinstatement test, Re-extinction: (b) | 10/10 | 20/20 | Reinstatement test + Re-extinction |
| Ehlers et al. | N=277 | Differential | Day 1: Acquisition Day 2: Extinction Renewal | 45.45% | Snowflakes | 1/1 | Electrotactile | Acquisition: (e) Extinction, Renewal: (a) | 11/11 | 23/23 | Renewal |

**Note.** The publications listed here do not constitute an exhaustive list of all studies on the research topic of interest. For a comprehensive overview on related studies, see pre-registered meta-analysis in preparation (Bruntsch et al., 2024). Detailed information on the included variables and reported results of the listed publications is provided in Supplementary Table 1.

[1] An avoidance phase took place between acquisition phase and extinction phase.

(a) No CS-US contingency information

(b) US information only (i.e., instructed that an US may occur, but not that its occurrence is related to the CS)

(c) General CS-US contingency information (i.e., instructed that the occurrence of the US is related to the CS, without information about the differential CS-US contingencies)

(d) Differential CS-US contingency information (i.e., instructed that one CS may be followed by the US whereas the other is not, without specifying which CS serves as CS+ and CS−)

(e) CS-specific contingency instruction (i.e., instructed about the contingency between the specific CS and the US, but not about the reinforcement rate)

n/a: not reported

**Supplementary Table 3.** Pearson correlation coefficients between individual anxiety-related trait scores (STAI-T, NEO-FFI-N, and IUS) and fear-conditioning outcome measures across experimental phases. Values are Pearson correlation coefficients (*r*) with Benjamini–Hochberg (BH)-adjusted *p*-values in parentheses. BH correction was applied separately for each anxiety-related trait measure across correlations with CS+, CS−, and CS discrimination. Significant correlations (*p*BH < .05) are shown in bold. Trend-level correlations (.05 ≤ *p*BH < .10) are shown in italics.

| **Measure** | **Phase** | **CS Discrimination** | | | **CS+** | | | **CS-** | | |
|---|---|---|---|---|---|---|---|---|---|---|
| | | STAI-T | NEO-FFI-N | IUS | STAI-T | NEO-FFI-N | IUS | STAI-T | NEO-FFI-N | IUS |
| SCR (raw) | Acquisition | .12 (.092) | .12 (.113) | .10 (.216) | .13 (.092) | .13 (.113) | .10 (.216) | .12 (.092) | .10 (.149) | .08 (.247) |
| | Full Extinction | .06 (.400) | .10 (.171) | .13 (.068) | .10 (.231) | .12 (.148) | .14 (.068) | .14 (.171) | .13 (.148) | .14 (.068) |
| | Early Extinction | .04 (.603) | .06 (.414) | .13 (.078) | .10 (.260) | .11 (.180) | .14 (.078) | .12 (.260) | .13 (.180) | .13 (.078) |
| | Late Extinction | .06 (.377) | .11 (140) | .11 (.124) | .10 (.221) | .12 (140) | .13 (.124) | .14 (.137) | .11 (140) | .12 (.124) |
| | Renewal | .10 (.173) | .11 (.178) | .09 (.346) | .10 (.173) | .11 (.178) | .09 (.346) | .10 (.173) | .09 (.212) | .06 (.436) |
| SCR (log, rc) | Acquisition | -.01 (.916) | .03 (.796) | .11 (.187) | .03 (.916) | .03 (.796) | .12 (.187) | .05 (.916) | .02 (.796) | .08 (.258) |
| | Full Extinction | .01 (.865) | .02 (.783) | .03 (.690) | .06 (.698) | .04 (.783) | .07 (.530) | .07 (.698) | .03 (.783) | .07 (.530) |
| | Early Extinction | .01 (.851) | .01 (.964) | .01 (.941) | .06 (.654) | .03 (.964) | .07 (.590) | .07 (.654) | .03 (.964) | .09 (.590) |
| | Late Extinction | .01 (.997) | .02 (.827) | .04 (.621) | .04 (.944) | .03 (.827) | .06 (.621) | .06 (.944) | .03 (.827) | .04 (.621) |
| | Renewal | -.01 (.938) | .10 (.463) | .02 (.824) | .05 (.827) | .09 (.463) | .05 (.824) | .07 (.827) | -.023 (.791) | .03 (.824) |
| FPS (raw) | Acquisition | .07 (.345) | .05 (.957) | -.04 (.548) | .09 (.345) | .02 (.957) | -.06 (.548) | .08 (.345) | -.01 (.957) | -.05 (.548) |
| | Full Extinction | .09 (.248) | .13 (.217) | .11 (.442) | .11 (.248) | .08 (.377) | .05 (.784) | .08 (.248) | .02 (.785) | -.01 (.873) |
| | Early Extinction | .07 (.302) | .13 (.237) | .09 (.707) | .11 (.299) | .10 (.286) | .05 (.793) | .09 (.299) | .04 (.572) | .01 (.963) |
| | Late Extinction | .12 (.152) | .14 (.153) | .12 (.322) | .12 (.152) | .08 (.444) | .05 (.591) | .06 (.419) | -.02 (.748) | -.04 (.591) |
| | Renewal | .17 (.051) | .13 (.428) | .10 (.428) | .09 (.329) | .02 (.958) | -.01 (.958) | -.01 (.912) | -.07 (.428) | -.08 (.428) |

| | | | | | | | | | | |
|---|---|---|---|---|---|---|---|---|---|---|
| FPS (T-score) | Acquisition | .09 (.326) | .11 (.176) | -.04 (.556) | .03 (.626) | .03 (.705) | -.13 (.217) | -.09 (.326) | -.13 (.176) | -.09 (.317) |
| | Full Extinction | .06 (.656) | .10 (.226) | .15 (.054) | .05 (.656) | .13 (.218) | .17 (.054) | -.03 (.656) | .03 (.723) | .01 (.957) |
| | Early Extinction | .08 (.422) | .15 (.056) | **.16 (.049)** | .08 (.422) | **.19 (.025)** | **.18 (.038)** | -.01 (.972) | .05 (.482) | .03 (.685) |
| | Late Extinction | .09 (.453) | .07 (.603) | .12 (.235) | .06 (.453) | .06 (.603) | .10 (.235) | -.05 (.453) | -.04 (.603) | -.04 (.592) |
| | Renewal | .10 (.239) | .03 (.654) | .06 (.580) | -.03 (.632) | -.08 (.424) | .02 (.798) | **-.18 (.033)** | -.14 (.160) | -.06 (.580) |
| US expectancy ratings | Acquisition | -.07 (.446) | -.05 (.609) | -.07 (.455) | .02 (.763) | -.01 (.993) | -.03 (.642) | .13 (.119) | .08 (.544) | .07 (.455) |
| | Full Extinction | .07 (.270) | .04 (.555) | -.01 (.986) | **.20 (.002)** | **.15 (.022)** | .07 (.414) | **.20 (.002)** | **.18 (.016)** | .10 (.295) |
| | Early Extinction | .04 (.547) | .02 (.708) | .01 (.890) | **.16 (.016)** | .13 (.054) | .08 (.317) | **.18 (.009)** | **.16 (.027)** | .11 (.255) |
| | Late Extinction | .10 (.105) | .05 (.436) | .01 (.978) | **.22 (.001)** | **.15 (.022)** | .06 (.481) | **.21 (.001)** | **.18 (.016)** | .10 (.391) |
| | Renewal | .07 (.265) | .03 (.615) | .01 (.908) | **.21 (.001)** | **.16 (.021)** | .05 (.595) | **.23 (.001)** | **.20 (.005)** | .07 (.595) |
| Fear ratings | Acquisition | .12 (.074) | .11 (.120) | .09 (.235) | .14 (.068) | **.16 (.042)** | .13 (.129) | .05 (.467) | .08 (.228) | .07 (.270) |
| | Extinction | **.13 (.039)** | **.16 (.015)** | .10 (.108) | **.19 (.006)** | **.22 (.001)** | **.20 (.004)** | **.16 (.018)** | **.13 (.045)** | **.19 (.004)** |
| | Renewal | .08 (.225) | .08 (.230) | -.01 (.927) | **.22 (.001)** | **.20 (.004)** | **.15 (.032)** | **.21 (.001)** | **.18 (.008)** | **.20 (.004)** |

**Note.** STAI-T = State-Trait Anxiety Inventory, Trait version; NEO-FFI-N = Neuroticism subscale of the NEO Five-Factor Inventory; IUS = Intolerance of Uncertainty Scale; SCR = skin conductance response; FPS = fear-potentiated startle; log = log-transformed; rc = range-corrected; sqrt = square-root-transformed; z = z-standardized.

**Supplementary Table 4.** Correlations between anxiety-related trait score and CS discriminations in several brain regions across phases

| Anatomical Region | Hemisphere | Voxels | T | *p* (< .001 | x | y | z |
|---|---|---|---|---|---|---|---|
| **Acquisition** | | | | | | | |
| *Positive correlation* | | | | | | | |
| Postcentral gyrus | L | 34 | 3.40 | < .001 | -57 | - | 32 |
| *Negative correlation* | | | | | | | |
| Anterior cingulate cortex | R | 18 | 3.75 | < .001 | 18 | 44 | 15 |
| Orbitofrontal gyrus | L | 17 | 3.52 | < .001 | -23 | 47 | -11 |
| **Early Extinction** | | | | | | | |
| *Positive correlation* | | | | | | | |
| Inferior temporal gyrus | L | 19 | 3.60 | < .001 | -42 | - | -8 |
| | R | 19 | 3.75 | < .001 | 44 | - | -10 |
| *Negative correlation* | | | | | | | |
| Ventral diencephalon | R | 38 | 4.13 | < .001 | 3 | - | -8 |
| **Late Extinction** | | | | | | | |
| *Negative correlation* | | | | | | | |
| Anterior cingulate cortex | R | 12 | 3.40 | < .001 | 4 | 38 | -4 |
| SMA/pre-SMA | L | 33 | 3.52 | < .001 | -36 | 26 | 30 |
| Anterior insula | R | 22 | 3.52 | < .001 | 39 | 21 | -8 |
| Thalamus | L | 10 | 3.39 | < .001 | -2 | - | 2 |
| Precentral gyrus | L | 17 | 3.45 | < .001 | -44 | -3 | 36 |
| Lingual gyrus | R | 32 | 3.66 | < .001 | 21 | - | -4 |
| | R | 14 | 3.36 | < .001 | 15 | - | 4 |
| Brain stem | R | 57 | 3.64 | < .001 | 14 | - | -10 |
| Cerebellum | R | 12 | 3.48 | < .001 | 8 | - | -42 |
| **Renewal** | | | | | | | |
| *Negative correlation* | | | | | | | |
| SMA/pre-SMA | L | 367 | 3.87 | < .001 | -6 | 24 | 48 |
| Posterior cingulate gyrus | R | 355 | 3.88 | < .001 | 6 | - | 20 |
| | R | 29 | 3.43 | < .001 | 10 | - | 0 |
| Anterior insula | L | 27 | 3.57 | < .001 | -38 | 8 | -14 |
| Thalamus | L | 15 | 3.56 | < .001 | -14 | - | 4 |
| Parahippocampal gyrus | L | 14 | 3.34 | < .001 | -15 | - | -12 |
| Superior temporal gyrus | R | 11 | 3.43 | < .001 | 62 | - | -3 |
| Middle temporal gyrus | L | 21 | 3.45 | < .001 | -57 | - | -2 |
| | L | 21 | 3.85 | < .001 | -46 | - | -15 |
| Superior frontal gyrus | L | 118 | 3.81 | < .001 | -4 | 57 | 15 |
| | L | 19 | 3.32 | < .001 | 21 | 32 | 52 |
| | R | 62 | 4.31 | < .001 | 12 | 38 | 22 |
| | R | 22 | 3.66 | < .001 | 21 | 48 | 39 |
| Middle frontal gryus | L | 139 | 3.96 | < .001 | -27 | 18 | 48 |
| | L | 15 | 3.34 | < .001 | -28 | 56 | 0 |
| | R | 38 | 3.43 | < .001 | 51 | 16 | 33 |
| | R | 31 | 3.36 | < .001 | 51 | 27 | 26 |
| Inferior frontal gyrus | L | 114 | 3.75 | < .001 | -56 | 14 | 12 |
| | R | 56 | 3.90 | < .001 | 51 | 30 | 12 |
| Orbitofrontal gyrus | L | 10 | 3.38 | < .001 | -27 | 20 | -24 |

**Note.** Voxel-wise threshold of $p < .001$ (uncorrected), minimum cluster size of k = 10 voxels.

#### Exploratory Factor Analyses

Currently, there is no consensus in this field of research regarding definitions and classifications of measures and latent dimensions of traits related to anxiety. In the past, the construct validity of the State-Trait Anxiety Inventory Trait version (STAI-T; Spielberger et al., 1983) has been criticized and terms like "Negative Emotionality" have been interpreted differently. For a deeper insight into this topic, see Klingelhöfer-Jens et al. (2025) and a pre-registered meta-analysis in preparation (Bruntsch et al., 2024). As the present study also investigates anxiety-related traits, a better understanding of the associations between the scale items and their latent dimensions is important. Thus, exploratory factor analyses (EFAs) were conducted. The aim was to examine content-related similarities between popular questionnaires in this field and to shed light on the constructs captured by the STAI-T and its relationship to other anxiety-related constructs.

In the present exploratory analyses, a first EFA was performed using data from the questionnaires employed in this study (EFA 1), followed by a second EFA that additionally included other questionnaires to allow more precise conclusions regarding the content represented by the resulting factors (EFA 2).

##### *EFA 1*

First, a parallel analysis of the STAI-T (Grimm, 2009; Spielberger et al., 1970), NEO-FFI-N (Gerhard, 1999; McCrae & Costa, 2004), and IUS (Buhr & Dugas, 2002; Gerlach et al., 2008) data was performed to explore the number of underlying factors. The suggested number of factors was five. In contrast, the scree test (see Supplementary Figure 1A) suggested two factors. Subsequently, EFAs were conducted using principal axis factoring as the extraction method. An oblique factor rotation (oblimin) was used because correlations between the factors were expected. Due to the differing recommendations regarding the number of factors, EFAs with two and three factors were calculated (see Supplementary Table 5). In total, the factors in the 3-Factor-Solution explained 40% of the variance. Of the explained variance, 51 % was accounted for by factor 1, 41% by factor 2, and 8% by factor 3. The factor intercorrelations indicated a moderate association between factors 1 and 2 ($r$ = .48) and weak associations between factor 3 and the other two factors ($r$ = .18 and $r$ = −.01). The model fit was acceptable (RMSR = .05; RMSEA = .053, 90% CI [0.049, 0.056]). Compared with the two-factor solution, the model fit improved when a third factor was included. However, a generally equivalent pattern emerged regardless of whether two or three factors were considered (see Supplementary Table 5): Items of the IUS loaded on a factor distinct from the STAI-T and NEO-FFI-N items, indicating that the IUS captures a conceptually distinct latent construct. In contrast, items from the STAI-T and NEO-FFI-N loaded on the same factor, suggesting that both scales reflect a common underlying latent dimension.

***EFA 2***

To obtain a more detailed picture of the classification and underlying latent structure, another exploratory factor analysis was conducted that included the STAI-T, NEO-FFI-N, the non-revised Anxiety Sensitivity Index (ASI; Alpers & Pauli, 2001; Reiss et al., 1986), and the Beck Depression Inventory II (BDI-II; Beck et al., 1996; Hautzinger et al., 2006). The Anxiety Sensitivity Index is a questionnaire designed to measure anxiety sensitivity, which refers to beliefs about the negative implications of experiencing anxiety. The ASI comprises 16 self-report items (e.g., “It scares me when I feel faint**”), which are rated on a five-point Likert scale ranging from 0 (very little) to 4 (very much). Total scores range from 0 to 64, with higher scores indicating greater anxiety sensitivity (Reiss et al., 1986). The Beck Depression Inventory II is a self-report measure intended to capture the severity of depressive symptoms during the past two weeks. The BDI-II includes 21 items (e.g., “Sadness”), each with four symptom-specific response options scored from 0 to 3, yielding sum scores between 0 and 63. Higher scores indicate more severe depressive symptoms (Beck et al., 1996).

The parallel analysis of the EFA 2 data suggested six factors, whereas the scree test suggested four factors (see Supplementary Figure 1B). Similar to EFA 1, principal axis factoring was used as the extraction method, together with an oblique factor rotation (oblimin). An EFA with four factors was calculated (see Supplementary Table 6). The four-factor solution explained a total of 38% of the variance. Of the explained variance, 44% was accounted for by factor 1, 24% by factor 2, 22% by factor 3, and 10% by factor 4. The factor intercorrelations indicated moderate associations between factors 1 and 2 ($r = .48$), factors 1 and 3 ($r = .57$), and factors 2 and 3 ($r = .33$), whereas the associations between factor 4 and the other factors were weak ($r = .28$, $r = .06$, and $r = .24$). The model fit was acceptable (RMSR = .05; RMSEA = .048, 90% CI [0.045, 0.051]).

Results showed that items from the NEO-FFI-N and most items from the STAI-T loaded on factor 1. Other STAI-T items loaded on factors 3 and 4. In contrast, ASI items mainly loaded on factor 2, whereas BDI-II items primarily loaded on factor 3 and partly on factor 4. Thus, the NEO-FFI-N, ASI, and BDI-II appeared to capture distinct latent dimensions. The STAI-T appeared to primarily measure the same latent construct as the NEO-FFI-N, while also showing similarities to the BDI-II and differing from the ASI. These findings suggest that the STAI-T may not primarily assess anxiety sensitivity but rather facets of depressive symptoms and broader negative emotionality, i.e., neuroticism.

**Supplementary Figure 1**

*Scree plots with parallel analysis and scree test results for EFA 1 (A) and EFA 2 (B)*

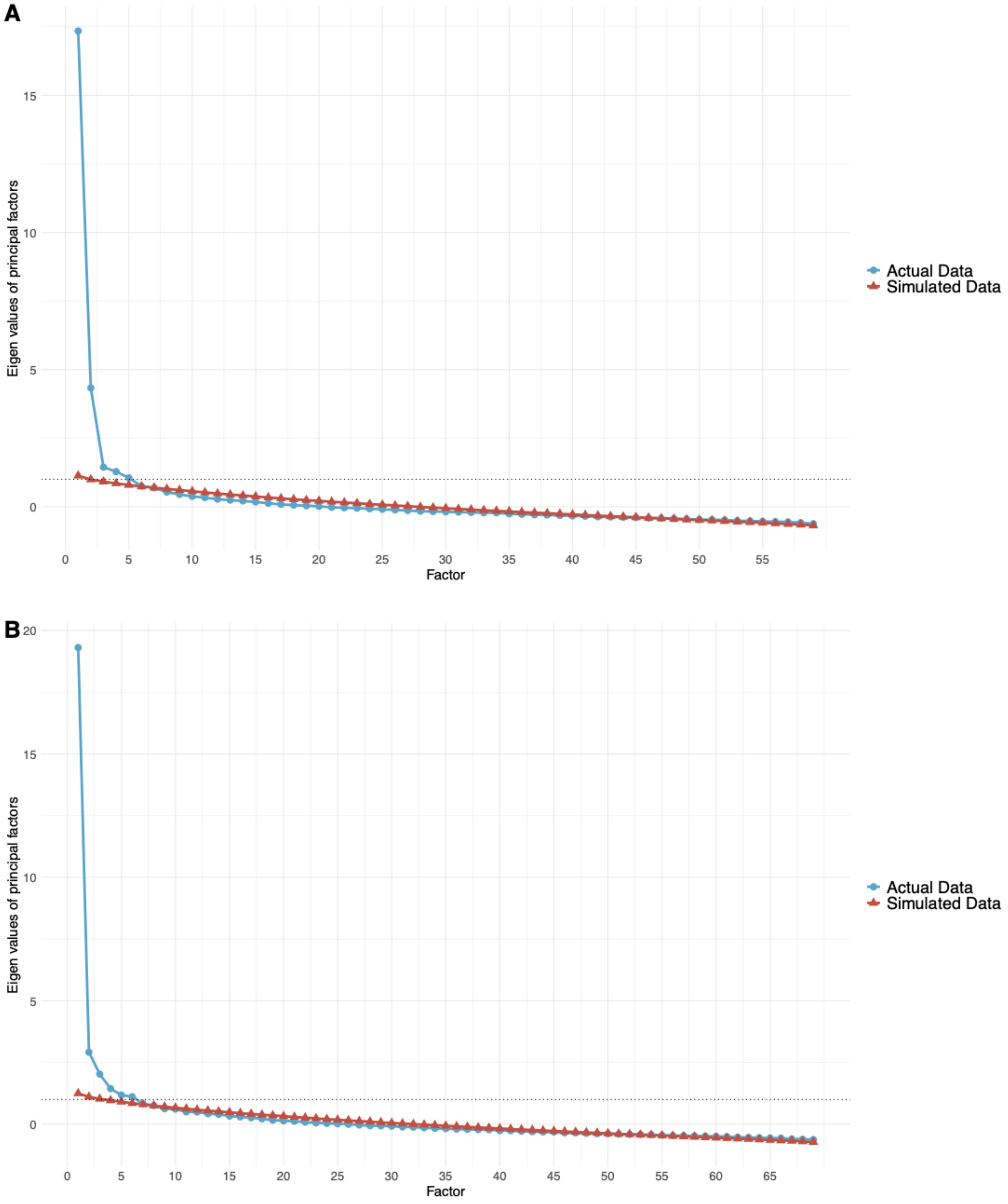

**Supplementary Table 5.** EFA 1 results using STAI-T, NEO-FFI-N, and IUS data.

| | | | | **2-Factor-Solution** | | **3-Factor-Solution** | | |
|---|---|---|---|---|---|---|---|---|
| **Scale** | **Item No** | **Items** | **Commu-nalities** | **Loadings F1** | **Loadings F2** | **Loadings F1** | **Loadings F2** | **Loadings F3** |
| STAI-T | 1 | I feel fine. | 0.40 | **0.53** | 0.06 | **0.39** | 0.16 | **0.35** |
| | 2 | I tire quickly. | 0.15 | **0.38** | -0.02 | **0.41** | -0.04 | -0.05 |
| | 3 | I feel like crying. | 0.41 | **0.67** | -0.11 | **0.70** | -0.13 | -0.04 |
| | 4 | I wish I could be as happy as others seems to be. | 0.43 | **0.66** | 0.00 | **0.62** | 0.03 | 0.11 |
| | 5 | I am losing opportunities because I cannot make decisions fast. | 0.25 | **0.34** | 0.19 | 0.24 | 0.25 | 0.22 |
| | 6 | I feel rested. | 0.31 | **0.56** | -0.04 | **0.51** | 0.00 | 0.14 |
| | 7 | I am calm. | 0.41 | **0.66** | -0.05 | **0.65** | -0.04 | 0.03 |
| | 8 | I feel that difficulties are piling up in such a way that I connot overcome them. | 0.61 | **0.71** | 0.12 | **0.66** | 0.15 | 0.13 |
| | 9 | I worry too much about things that do not really matter. | 0.43 | **0.51** | 0.08 | **0.65** | -0.02 | -0.30 |
| | 10 | I am happy. | 0.51 | **0.62** | 0.00 | **0.46** | 0.11 | **0.41** |
| | 11 | I am inclined to take things hard. | 0.46 | **0.57** | 0.18 | **0.58** | 0.17 | -0.01 |
| | 12 | I lack self-confidence. | 0.42 | **0.59** | 0.11 | **0.55** | 0.13 | 0.08 |
| | 13 | I feel secure. | 0.39 | **0.50** | 0.06 | **0.35** | 0.16 | **0.37** |
| | 14 | I try to avoid facing a crisis or difficulty. | 0.35 | **0.52** | 0.07 | **0.58** | 0.03 | -0.15 |
| | 15 | I feel blue. | 0.50 | **0.76** | -0.14 | **0.71** | -0.10 | 0.14 |
| | 16 | I am content. | 0.51 | **0.61** | -0.01 | **0.43** | 0.11 | **0.45** |
| | 17 | Some unimportant thoughts run through my mind and bother me. | 0.46 | **0.59** | 0.04 | **0.70** | -0.04 | -0.23 |
| | 18 | I take disappointsments so keenly that I connot get them out of my mind. | 0.38 | **0.51** | 0.15 | **0.56** | 0.12 | -0.11 |
| | 19 | I am a steady person. | 0.53 | **0.75** | -0.07 | **0.70** | -0.03 | 0.15 |
| | 20 | I become tense and upset when I think about my current concerns. | 0.48 | **0.62** | 0.07 | **0.68** | 0.03 | -0.13 |
| NEO-FFI-N | 1 | I am not a worrier. | 0.31 | **0.49** | 0.11 | **0.52** | 0.09 | -0.07 |
| | 2 | I often feel inferior to others. | 0.37 | **0.52** | 0.15 | **0.51** | 0.16 | -0.02 |
| | 3 | When I'm under a great deal of stress, sometimes I feel like I'm going to pieces. | 0.44 | **0.63** | 0.03 | **0.67** | 0.01 | -0.07 |
| | 4 | I rarely feel lonely or blue. | 0.44 | **0.67** | -0.02 | **0.62** | 0.02 | 0.13 |
| | 5 | I often feel tense and jittery. | 0.56 | **0.71** | 0.06 | **0.73** | 0.05 | -0.03 |
| | 6 | Sometimes I feel completely worthless. | 0.50 | **0.69** | 0.03 | **0.67** | 0.05 | 0.07 |
| | 7 | I rarely feel fearful or anxious. | 0.35 | **0.59** | -0.04 | **0.63** | -0.06 | -0.06 |
| | 8 | I often get angry at the way people treat me. | 0.22 | **0.39** | 0.13 | **0.40** | 0.13 | -0.01 |
| | 9 | Too often, when things go wrong, I get discouraged and feel like giving up. | 0.31 | **0.44** | 0.18 | **0.40** | 0.21 | 0.08 |
| | 10 | I am seldom sad or depressed. | 0.55 | **0.76** | -0.05 | **0.71** | -0.01 | 0.13 |
| | 11 | I often feel helpless and want someone else to solve my problems. | 0.39 | **0.51** | 0.18 | **0.48** | 0.21 | 0.09 |
| | 12 | At times I have been so ashamed I just wanted to hide. | 0.26 | **0.32** | 0.24 | **0.39** | 0.19 | -0.17 |
| IUS | 1 | Uncertainty stops me from having a firm opinion. | 0.24 | 0.24 | **0.33** | 0.23 | **0.34** | 0.02 |
| | 2 | Being uncertain means that a person is disorganized. | 0.23 | 0.01 | **0.42** | -0.09 | **0.48** | 0.20 |
| | 3 | Uncertainty makes life intolerable. | 0.40 | 0.00 | **0.62** | -0.04 | **0.65** | 0.05 |

| | | | | | | | |
|---|---|---|---|---|---|---|---|
| 4 | It's unfair not having any guarantees in life. | 0.18 | 0.00 | **0.43** | 0.03 | **0.41** | -0.08 |
| 5 | My mind can't be relaxed if I don't know what will happen tomorrow. | 0.36 | 0.08 | **0.47** | 0.21 | **0.38** | **-0.33** |
| 6 | Uncertainty makes me uneasy, anxious, or stressed. | 0.45 | **0.31** | **0.41** | **0.42** | **0.34** | -0.25 |
| 7 | Unforeseen events upset me greatly. | 0.52 | 0.16 | **0.60** | 0.24 | **0.54** | -0.23 |
| 8 | It frustrates me not having all the information I need. | 0.37 | 0.06 | **0.51** | 0.17 | **0.44** | **-0.29** |
| 9 | Uncertainty keeps me from living a full life. | 0.51 | 0.13 | **0.64** | 0.10 | **0.66** | 0.04 |
| 10 | One should always look ahead so as to avoid surprises. | 0.32 | -0.21 | **0.61** | -0.13 | **0.55** | -0.23 |
| 11 | A small unforeseen event can spoil everything, even with the best of planning. | 0.28 | -0.08 | **0.56** | -0.11 | **0.58** | 0.03 |
| 12 | When it's time to act, uncertainty paralyses me. | 0.57 | 0.12 | **0.66** | 0.04 | **0.72** | 0.16 |
| 13 | Being uncertain means that I am not first rate. | 0.34 | -0.02 | **0.57** | -0.08 | **0.61** | 0.11 |
| 14 | When I am uncertain, I can't go forward. | 0.58 | 0.18 | **0.62** | 0.07 | **0.69** | 0.22 |
| 15 | When I am uncertain I can't function very well. | 0.45 | 0.16 | **0.58** | 0.12 | **0.60** | 0.06 |
| 16 | Unlike me, others always seem to know where they are going with their lives. | 0.39 | 0.17 | **0.51** | 0.11 | **0.55** | 0.12 |
| 17 | Uncertainty makes me vulnerable, unhappy, or sad. | 0.58 | 0.21 | **0.64** | 0.18 | **0.66** | 0.05 |
| 18 | I always want to know what the future has in store for me. | 0.41 | **-0.35** | **0.71** | -0.27 | **0.65** | -0.25 |
| 19 | I can't stand being taken by surprise. | 0.28 | -0.07 | **0.56** | -0.06 | **0.55** | -0.04 |
| 20 | The smallest doubt can stop me from acting. | 0.33 | 0.17 | **0.47** | 0.14 | **0.49** | 0.07 |
| 21 | I should be able to organize everything in advance. | 0.42 | -0.12 | **0.68** | -0.05 | **0.63** | -0.20 |
| 22 | Being uncertain means that I lack confidence. | 0.35 | 0.24 | **0.43** | 0.18 | **0.47** | 0.12 |
| 23 | I think it's unfair that other people seem sure about their future. | 0.34 | 0.01 | **0.55** | -0.05 | **0.59** | 0.13 |
| 24 | Uncertainty keeps me from sleeping soundly. | 0.27 | 0.06 | **0.46** | 0.13 | **0.42** | -0.18 |
| 25 | I must get away from all uncertain situations. | 0.34 | -0.01 | **0.59** | -0.02 | **0.59** | 0.00 |
| 26 | The ambiguities in life stress me. | 0.52 | 0.19 | **0.61** | 0.22 | **0.59** | -0.09 |
| 27 | I can't stand being undecided about my future. | 0.49 | -0.05 | **0.73** | -0.03 | **0.71** | -0.09 |

**Note.** Communalities of the 3-Factor-Solution are shown. Standardized loadings are based on correlation matrix. Loadings higher than or equal to .3 are formatted in bold. F = Factor.

**Supplementary Table 6.** EFA 2 results using STAI-T, NEO-FFI-N, ASI, and BDI-II data.

| | | | | 4-Factor-Solution | | | |
|---|---|---|---|---|---|---|---|
| **Scale** | **Item No** | **Items** | **Commu-nalities** | **Loadings F1** | **Loadings F2** | **Loadings F3** | **Loadings F4** |
| STAI-T | 1 | I feel fine. | .40 | **0.54** | -0.06 | -0.03 | 0.27 |
| | 2 | I tire quickly. | .29 | 0.19 | 0.06 | **0.38** | -0.28 |
| | 3 | I feel like crying. | .45 | **0.36** | 0.10 | **0.35** | -0.08 |
| | 4 | I wish I could be as happy as others seems to be. | .47 | **0.39** | 0.09 | 0.25 | 0.18 |
| | 5 | I am losing opportunities because I cannot make decisions fast. | .24 | 0.24 | 0.23 | 0.02 | 0.19 |
| | 6 | I feel rested. | .33 | **0.52** | -0.10 | 0.16 | -0.02 |
| | 7 | I am calm. | .42 | **0.64** | 0.05 | -0.01 | -0.05 |
| | 8 | I feel that difficulties are piling up in such a way that I connot overcome them. | .58 | **0.60** | 0.14 | 0.12 | 0.02 |
| | 9 | I worry too much about things that do not really matter. | .36 | **0.46** | 0.12 | 0.14 | -0.26 |
| | 10 | I am happy. | .49 | **0.57** | -0.04 | -0.02 | **0.31** |
| | 11 | I am inclined to take things hard. | .48 | **0.42** | 0.26 | 0.14 | 0.05 |
| | 12 | I lack self-confidence. | .44 | **0.60** | 0.12 | -0.07 | 0.12 |
| | 13 | I feel secure. | .41 | **0.54** | -0.15 | -0.01 | **0.31** |
| | 14 | I try to avoid facing a crisis or diffi-culty. | .35 | **0.44** | 0.19 | 0.08 | -0.09 |
| | 15 | I feel blue. | .52 | **0.40** | 0.16 | 0.22 | 0.18 |
| | 16 | I am content. | .47 | **0.61** | -0.14 | -0.03 | 0.29 |
| | 17 | Some unimportant thoughts run through my mind and bother me. | .39 | **0.48** | 0.19 | 0.09 | -0.12 |
| | 18 | I take disappointsments so keenly that I connot get them out of my mind. | .38 | **0.45** | 0.24 | 0.03 | -0.01 |
| | 19 | I am a steady person. | .53 | **0.73** | -0.09 | 0.08 | -0.03 |
| | 20 | I become tense and upset when I think about my current concerns. | .48 | **0.65** | -0.01 | 0.13 | -0.18 |
| NEO-FFI-N | 1 | I am not a worrier. | .34 | **0.57** | 0.12 | -0.06 | -0.11 |
| | 2 | I often feel inferior to others. | .39 | **0.59** | 0.13 | -0.11 | 0.07 |
| | 3 | When I'm under a great deal of stress, sometimes I feel like I'm going to pieces. | .45 | **0.61** | 0.01 | 0.13 | -0.16 |
| | 4 | I rarely feel lonely or blue. | .44 | **0.57** | 0.02 | 0.09 | 0.08 |
| | 5 | I often feel tense and jittery. | .57 | **0.73** | 0.02 | 0.06 | -0.12 |
| | 6 | Sometimes I feel completely worthless. | .52 | **0.62** | 0.12 | -0.01 | 0.11 |
| | 7 | I rarely feel fearful or anxious. | .35 | **0.58** | 0.07 | -0.02 | -0.09 |
| | 8 | I often get angry at the way people treat me. | .22 | **0.38** | 0.06 | 0.07 | 0.04 |
| | 9 | Too often, when things go wrong, I get discouraged and feel like giving up. | .33 | **0.57** | 0.12 | -0.16 | 0.03 |
| | 10 | I am seldom sad or depressed. | .54 | **0.68** | 0.02 | 0.07 | 0.01 |
| | 11 | I often feel helpless and want someone else to solve my problems. | .37 | **0.53** | 0.05 | 0.08 | 0.01 |
| | 12 | At times I have been so ashamed I just wanted to hide. | .23 | **0.43** | 0.15 | -0.06 | -0.06 |
| ASI | 1 | It is important to me not to appear nervous. | .14 | 0.12 | 0.22 | -0.06 | -0.23 |
| | 2 | When I cannot keep my mind on a task, I worry that I might be going crazy. | .28 | -0.03 | **0.47** | 0.14 | 0.07 |
| | 3 | It scares me when I feel „shaky“ (trembling). | .55 | 0.08 | **0.73** | -0.10 | -0.08 |
| | 4 | It scares me when I feel faint | .45 | 0.09 | **0.63** | -0.05 | 0.07 |

| | | | | | | | |
|---|---|---|---|---|---|---|---|
| | 5 | It is important to me to stay in control of my emotions. | .17 | 0.04 | 0.19 | -0.03 | **0.34** |
| | 6 | It scares me when my heart beats rapidly. | .36 | 0.06 | **0.57** | 0.01 | -0.13 |
| | 7 | It embarrasses me when my stomach growls. | .19 | 0.22 | 0.22 | 0.11 | -0.14 |
| | 8 | It scares me when I am nauseous. | .32 | 0.00 | **0.58** | -0.03 | -0.05 |
| | 9 | When I notice that my heart is beating rapidly, I worry that I might have a heart attack. | .16 | -0.03 | **0.36** | 0.12 | 0.02 |
| | 10 | It scares me when I become short of breath. | .27 | -0.05 | **0.55** | -0.03 | 0.00 |
| | 11 | When my stomach is upset, I worry that I might bc seriously ill. | .25 | -0.08 | **0.51** | 0.05 | 0.02 |
| | 12 | It scares we when I am unable to keep my mind on a task. | .36 | -0.02 | **0.61** | -0.07 | 0.11 |
| | 13 | Other people notice when I feel shaky. | .18 | 0.10 | **0.35** | 0.03 | -0.10 |
| | 14 | Unusual body sensations scare me. | .41 | 0.05 | **0.57** | 0.12 | -0.10 |
| | 15 | When I am nervous, I worry that I might be mentally ill. | .41 | 0.07 | **0.47** | 0.20 | 0.11 |
| | 16 | It scams me when I am nervous. | .46 | -0.03 | **0.66** | 0.06 | 0.10 |
| BDI-II | 1 | Sadness | .48 | 0.26 | 0.01 | **0.35** | 0.29 |
| | 2 | Pessimism | .28 | 0.23 | 0.10 | 0.15 | 0.25 |
| | 3 | Past Failure | .39 | **0.34** | 0.05 | 0.22 | 0.20 |
| | 4 | Loss of Pleasure | .38 | 0.12 | 0.02 | **0.45** | 0.20 |
| | 5 | Guilty Feelings | .37 | 0.21 | 0.11 | **0.36** | 0.12 |
| | 6 | Punishment Feelings | .36 | -0.01 | 0.17 | **0.31** | **0.38** |
| | 7 | Self-Dislike | .45 | 0.29 | -0.06 | 0.27 | **0.36** |
| | 8 | Self-Criticalness | .33 | 0.19 | 0.14 | **0.30** | 0.15 |
| | 9 | Suicidal Thoughts or Wishes | .34 | -0.07 | 0.23 | **0.32** | **0.36** |
| | 10 | Crying | .38 | -0.06 | -0.05 | **0.66** | 0.02 |
| | 11 | Agitation | .46 | -0.01 | 0.01 | **0.70** | -0.09 |
| | 12 | Loss of Interest | .46 | 0.10 | 0.09 | **0.41** | **0.33** |
| | 13 | Indecisiveness | .36 | 0.21 | 0.15 | **0.32** | 0.13 |
| | 14 | Worthlessness | .45 | 0.21 | 0.06 | 0.26 | **0.39** |
| | 15 | Loss of Energy | .45 | 0.13 | -0.01 | **0.61** | -0.11 |
| | 16 | Changes in Sleeping Pattern | .25 | -0.10 | 0.07 | **0.54** | -0.05 |
| | 17 | Irritability | .38 | 0.01 | 0.02 | **0.59** | 0.04 |
| | 18 | Changes in Appetite | .35 | -0.04 | 0.01 | **0.60** | 0.05 |
| | 19 | Concentration Difficulty | .50 | 0.14 | -0.02 | **0.63** | -0.05 |
| | 20 | Tiredness or Fatigue | .21 | -0.01 | 0.04 | 0.21 | **0.35** |
| | 21 | Loss of Interest in Sex | .44 | 0.07 | 0.07 | **0.53** | 0.16 |

**Note.** Communalities of the 4-Factor-Solution are shown. Standardized loadings are based on correlation matrix. Loadings higher than or equal to .3 are formatted in bold. F = Factor.
